# Farnesyl Transferase Inhibition for the Treatment of Tauopathies

**DOI:** 10.1101/500801

**Authors:** Israel Hernandez, Gabriel Luna, Jennifer N. Rauch, Michel Giroux, Celeste M. Karch, Daniel Boctor, Nadia J. Storm, Antonio Diaz, Cezary Zekanowski, Alexander A. Kang, Cassidy Hinman, Vesna Cerovac, Elmer Guzman, Honjun Zhou, Alison Goate, Steven K. Fisher, Ana M. Cuervo, Kenneth S. Kosik

## Abstract

Tau inclusions are a shared feature of many neurodegenerative conditions and tau mutations lead to frontotemporal dementia. Approaches to treatment of these conditions have focused directly on the tau protein by targeting its post-translational modifications, its levels and its tendency to aggregate. We discovered a novel regulatory pathway for tau degradation that operates through the Rhes protein, a GTPase. Rhes is farnesylated and treatment with the farnesyl transferase inhibitor, lonafarnib, reduced Rhes, attenuated behavioral abnormalities, significantly reduced atrophy, tau inclusions, sumoylation and ubiquitination, as well as microgliosis in the rTg4510 tauopathy mouse. Direct reduction of Rhes levels reproduced the results observed with lonafarnib. The mechanism of lonafarnib action, as mediated by Rhes to reduce tau pathology, operates through the lysosome without involvement of the proteasome. Finally we show that the developmental increase in Rhes levels can be homeostatically regulated in the presence of tau mutations as a protective mechanism through which cells sense abnormal tau before any pathology is present. The extensive human trials of lonafarnib for other conditions, makes this drug ideal for repurposing to treat tauopathies.

**One-sentence summary:** Via a mechanism that involves targeting Rhes, lonafarnib can induce lysosomal-mediated tau degradation and prevent pathology in a tau mouse model

## Introduction

The tauopathies constitute a broad range of diseases all of which share the hallmark feature of tau inclusions. Although tau-related diseases including Alzheimer’s disease and chronic traumatic encephalopathy are serious public health problems (1), no disease-modifying treatment currently exists for these conditions. Generally, approaches to treatment have directly targeted the tau protein and include tau antibodies (2), antisense oligonucleotides (3), caspase cleavage products (4), and anti-aggregation agents (5); however, few pharmacologic interventions directed toward tau pathways have reached clinical trials. Interventions in upstream pathways such as tau phosphorylation (6) and acetylation inhibitors (7) have shown some efficacy in animal models; however, none of these approaches has had success in human clinical trials. Rapidly growing interest in autophagy as a downstream pathway that mediates tau clearance (8) as well as the implication of autophagy and lysosomes in other neurodegenerative conditions (9–11) suggests therapeutic opportunities. One potentially druggable pathway linked to autophagy has been suggested in Huntington’s disease (HD). This pathway is mediated by the *RASD2* gene, which encodes the Rhes protein, a small GTPase member of the Ras superfamily (12). *RASD2* became of interest in HD because it was thought to be expressed mainly in the striatum (13); however, in humans, Rhes expression is clearly evident in the cerebral cortex (14). Rhes activates autophagy independently of mTOR via interaction with Beclin 1 (15). Furthermore, Rhes has been shown to modulate the aggregation state of mutant Huntingtin (mHtt) by promoting its sumoylation, thus reducing cell survival (16). This toxicity involves binding of Rhes to mHtt and requires the Rhes CXXX farnesylated membrane attachment site (16). We became interested in these findings because the induction of autophagy is considered a pharmacological class effect of farnesyl transferase inhibition (17) and therefore could have broader effects than those mediated by the Rhes pathway alone and be relevant for tauopathies. Therefore we studied farnesyl transferase as a therapeutic target for the treatment of tauopathies and present the use of the farnesyl transferase inhibitor, lonafarnib, a drug with which there is extensive clinical experience (18–20), as a proof of concept.

## Results

### Farnesylation inhibition attenuates tau pathology in rTg4510 mice

Lonafarnib is a potent farnesyl transferase inhibitor with a K_i_ in the nM range (21), crosses the blood brain barrier and has few minor side effects in humans. We sought to determine the effects of lonafarnib administration in the rTg4510 mouse, a widely used model for frontotemporal dementia (22). These mice were reported to develop tau tangles in the cerebral cortex by 4 months of age and in the hippocampus by 5.5 months. By 5.5 months, the mice also lose about 60% of their hippocampal CA1 neurons (22). Spatial memory deficits become apparent by 2.5 to 4 months of age; however electrophysiological properties of cortical neurons are affected before the accumulation of tau pathology (23, 24). To determine precisely the effects of lonafarnib we replicated the time course for the onset of pathology in rTg4510 (Fig. 1; supplemental Fig. 1). Similar to the reported data, MC1 immunoreactivity, corresponding to neurofibrillary tangles throughout hippocampus, amygdala, entorhinal cortex and cerebrocortex (Fig. 1A; green), was first detected at ~16 weeks (supplemental Fig. 1) and progressed by 20 weeks to a mean of 92.99±13.63 MC1^+^ cells/mm^2^ in the cerebral cortex, and 67.57±13.11 cells/mm^2^ in the hippocampus (Fig. 1D, E). Wild type (WT) mice showed no detectable MC1^+^ neurons (Fig. 1C).

**Figure 1.**
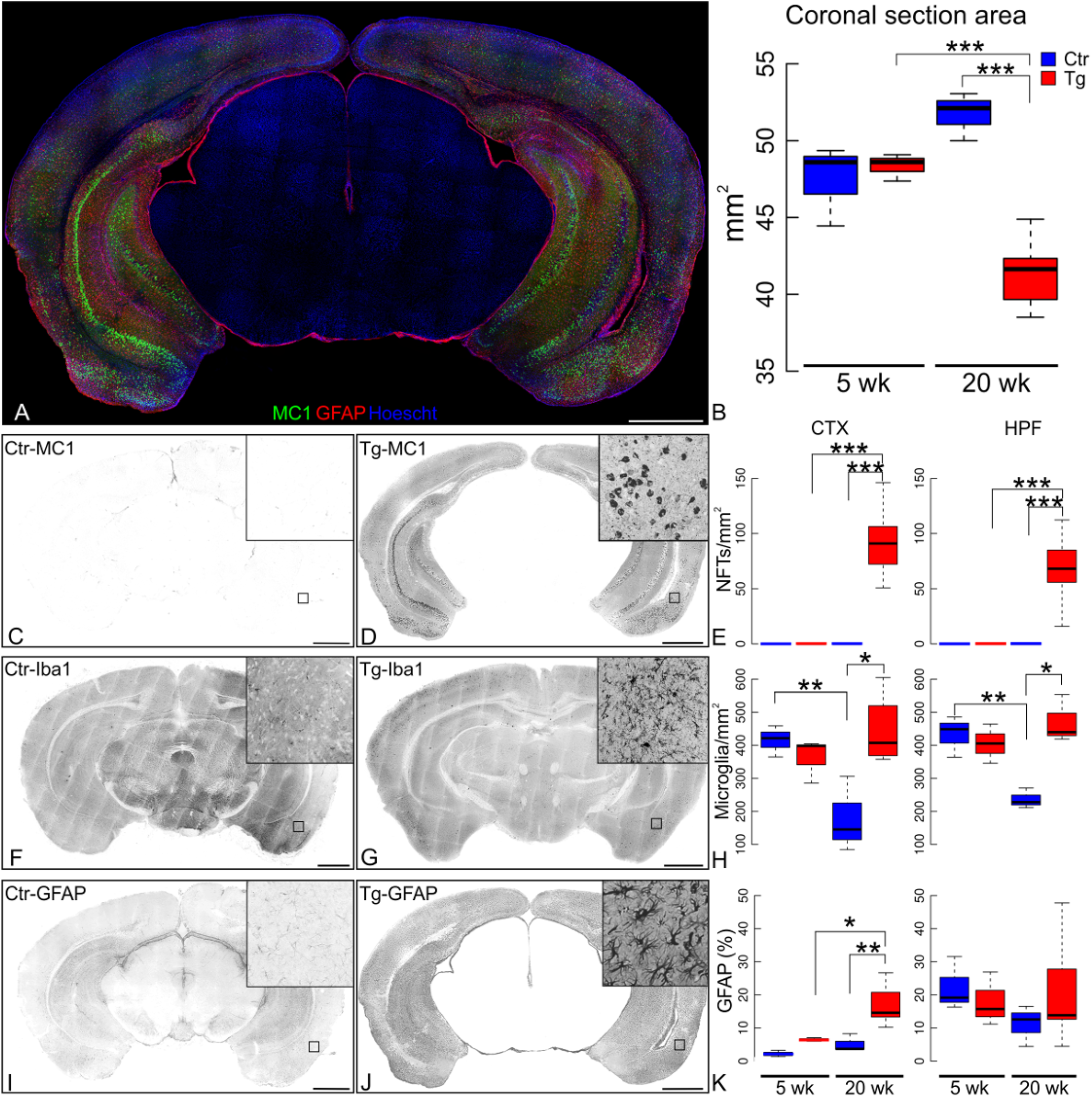
Baseline time course of pathology in the rTg4510 mouse model. (A) Representative image of brain coronal section tiling 800 x 800 pixels Z-Stacks mosaic labeled with MC1 (Green), GFAP (Red) and Hoechst (cell nuclei, blue) is shown. (B) The coronal section area of the micrographs was computed. At five weeks, rTg4510 transgenic mice (Tg) did not differ in brain size from their non-transgenic littermates (p=0.942), however at 20 weeks, the area was significantly reduced in transgenic mice compared to both age-matched controls and younger transgenic (p=6.94×10^−4^). (C) 20-week-old control littermates (Ctr) are non-reactive to MC1 staining; (D) whereas 20 week old transgenic mice have a high density of strongly labeled neurons in both cerebral cortex (CTX) and hippocampal formation (HPF). (E) Quantification of NFTs/mm^2^ in CTX and HPF. Representative micrographs showing microglia (Iba1^+^ cells) in 20 week old Ctr (F) and Tg (G) mice, and (H) quantification of microglia/mm^2^ density, revealing that microglia lacks an age-related decline in rTg4510 transgenic mice. At 20-weeks, in the Ctr CTX there were 178.45±66.19 microglia/mm^2^ and in the Tg CTX there were 444.20±55.91 microglia/mm^2^ (p=1.9×10^−3^). In the Ctr HPF, there were 236.74±17.68 microglia/mm^2^ and in the Tg HPF there were 470.99±41.97 microglia/mm^2^ (p=5.11×10^−3^). (I) Astrocytes labeled with GFAP antibody in non-transgenic (Ctr) and (J) 20 week-old transgenic (Tg) mice show cortical astrogliosis accompanying the MC1 immunoreactivity in the rTg4510 mouse model. Activated hypertrophied astrocytes are shown in the inset. (K) GFAP^+^ percentage area in the coronal sections was quantified. The cortical GFAP signal (ANOVA p=9.14×10^−4^) quantified in Tg at 20 weeks increased in comparison of age-matched Ctr (p=9.2×10^−3^). The cortical GFAP signal also increased in 20-week-old transgenic (Tg) compared to five-week-old Tg (p=0.020). The hippocampal GFAP signal between Tg and Ctr mice at 5 and 20 weeks did not differ (ANOVA p=0.676). Statistics shown for post-hoc Tukey-HSD test at *p<0.05, **p<0.01, ***p<1×10^−3^. Scale bar scale 1 mm.

Lonafarnib treatment was initiated at ten weeks of age. Lonafarnib was resuspended in 20% 2-Hydroxypropyl)-β-cyclodextrin as vehicle and gavage fed on an intermittent schedule, five days on five days off at 80 mg/Kg/day (25). The drug was remarkably effective in reducing MC1 immunoreactivity. When evaluated at 20-weeks (Fig. 2B-H), the MC1 immunoreactivity in the lonafarnib treated cortex (44.27±4.064 MC1^+^ cells/mm^2^) and hippocampus (13.38±2.615 MC1+ cells/mm^2^) was significantly reduced compared to either vehicle-treated (Fig. 2H; 84.15±5.050 cells/mm^2^ in cortex and 42.01±3.305 cells/mm^2^ in hippocampus; cortex p=7.2×10^−3^, hippocampus p=6.4×10^−3^), or untreated age-matched transgenic mice (cortex p=3.5×10^−3^, hippocampus p=2.4×10^−5^). In contrast to chronic administration, lonafarnib was ineffective as an acute intervention. For this experiment, animals were gavage-treated daily for 2 weeks (80 mg/Kg/day) in 20 week-old rTg4510 mice, after tau pathology was widespread. This treatment failed to alter the location and the extent of the MC1 pathology observed when compared to age-matched vehicle treated and untreated transgenic mice (data not shown).

**Figure 2.**
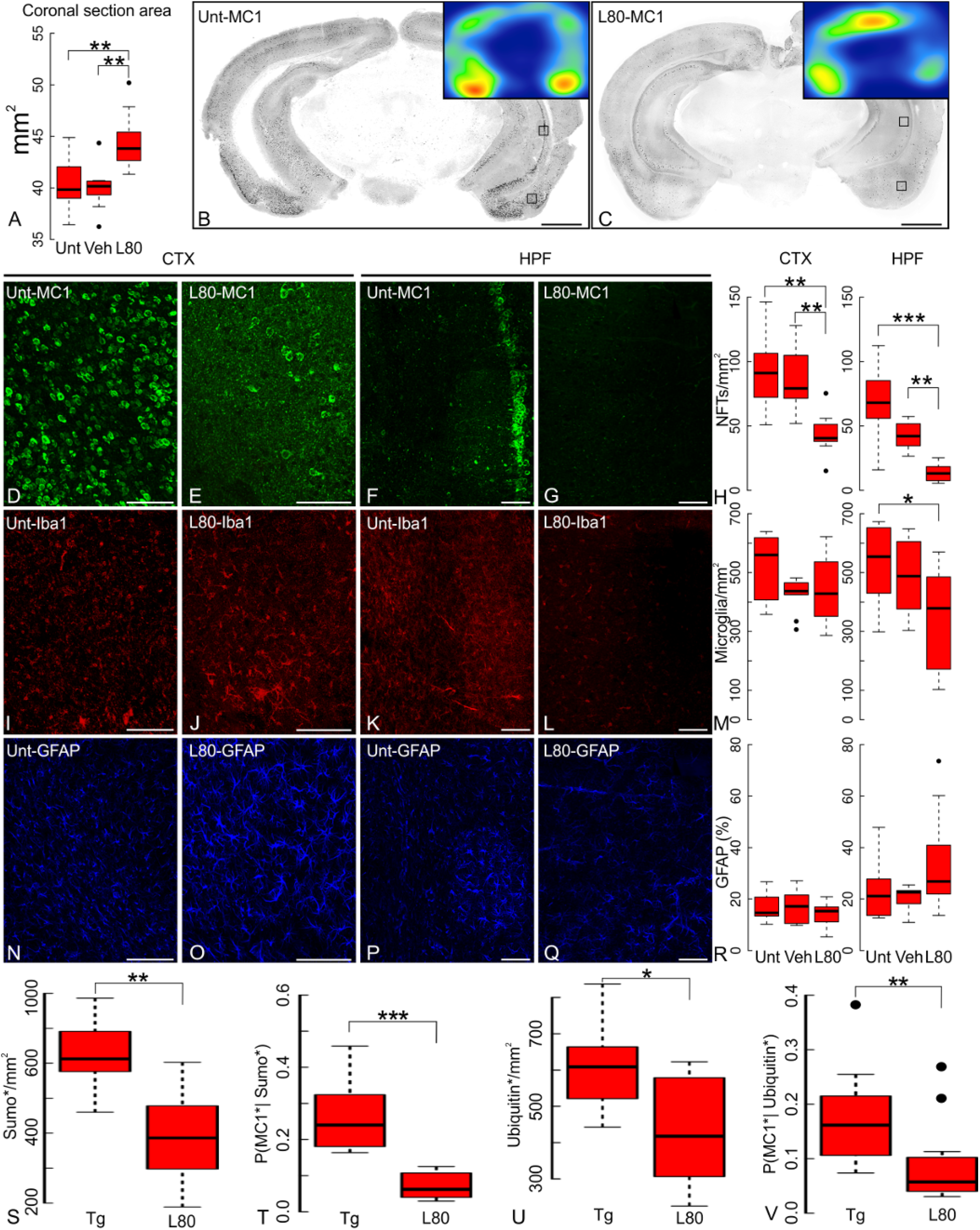
Lonafarnib treatment prevents neurofibrillary tangle formation and microgliosis. (A) Brain coronal section area in 20-week old rTg4510 transgenic mice (Tg) that received chronic oral administration of Lonafarnib (L80) is larger than that of untreated (Unt) or vehicle alone (Veh) Tg mice. Untreated and vehicle-treated mice measured 40.43±0.83 mm^2^ and 40.02±0.73 mm^2^, respectively. Coronal section areas from lonafarnib-treated Tg animals averaged 44.55±0.95 mm^2^ (untreated p=5.6×10^−3^, vehicle-treated p=2.4×10^−3^). (B-C) Reduction of the extent of MC1 immunoreactivity in lonafarnib (L80) treated transgenic mice compared to untreated mice (Unt). Scale bar 1 mm. (D-G) Detail of insets of panels B and C showing representative MC1 immunoreactivity on cortex (CTX) and hippocampus (HPF) of either untreated (Unt) or lonafarnib treated (L80) in 20 week-old Tg mice; (H) large-scale coronal section mosaics quantified for MC1^+^/mm^2^ indicates a significant reduction of tau pahology after lonafarnib treatment (L80) when compared to untreated animals (Unt) or animals treated with vehicle alone (Veh). (I-L) Density of microglia in the CTX and HPF of transgenic mice treated with lonafarnib (L80) or left untreated (Unt) is shown by Iba1 immunolabeling. Hippocampal microglial reactivity declined upon lonafarnib treatment. (M) Microglia quantification of coronal section mosaics in both CTX and HPF of transgenic mice treated accordingly. In the HPF (ANOVA p=0.040), lonafarnib-treated Tg mice at 20 weeks had 343.33±13.24 Iba1^+^ cells/mm^2^ compared to vehicle treated animals (488.38±11.53 Iba1+ cells/mm^2^) or untreated controls (526.95±12.02 Iba1^+^ cells/mm^2^, p=0.05) and represents a significant reduction in microgliosis. No statistically significant differences were observed in the CTX (ANOVA p=0.211) of lonafarnib-treated animals (442.51±10.73 Iba1^+^ cells/mm^2^) when compared to vehicle (425.30±7.94 Iba1^+^ cells/mm^2^) or untreated animals (515.45±11.03 Iba1^+^ cells/mm^2^). (N-Q) Astrocytes immunostained with GFAP in CTX or HPF of untreated and lonafarnib treated Tg mice, and (R) quantification of GFAP signal in full coronal slices. Neither lonafarnib (L80) nor vehicle alone (Veh) altered astrocytes in 20 weeks-old Tg mice (cortex ANOVA p=0.411, hippocampus ANOVA p=0.111). Statistics shown for post-hoc Tukey-HSD test at *p<0.05, **p<0.01, ***p<1×10^−3^. (S) Lonafarnib treatment (L80) reduced sumoylated cells compared to rTg4510 mice (Tg; p=2.55×10^−3^). (T) Lonafarnib reduced ubiquitin labeled cells compared to rTg4510 mice (Tg; p=0.05). Counts of double-labeled cells in the micrographs were used to compute Bayesian probabilities of any cell to be tau immunoreactive (MC1^+^) given sumo^+^ (U) or ubiquitin^+^ (V) immunoreactivity. Lonafarnib also decreased these probabilities of double-labeling, in comparison to age-matched rTg4510 mice (Tg). Statistics shown for post-hoc Tukey-HSD test at *p<0.05, **p<0.01, ***p<1×10^−3^.Scale bar 100 μm.

Chronic lonafarnib treatment also prevented the reduction in brain size among aged transgenic mice. Coronal section area of littermate controls (Ctr) and transgenic (Tg) mice did not differ at five weeks, but significantly diminished by 20 weeks, as mice aged (Fig. 1B). Lonafarnib-treated Tg (L80) mice at 20-weeks had significantly larger coronal brain areas than both vehicle (Veh) and Untreated (Unt) age-matched transgenic mice (Fig. 2A; ANOVA p=1.32×10^−3^).

Microglia counts, assessed by Iba1 at 5 and 20 weeks (Fig. 1F-G; supplemental Fig. 1), decreased significantly with age (Fig. 1H) in Ctr cortex (p=0.049) and hippocampus (p=0.014), but remained elevated in 20 week old Tg mice. At five weeks, no statistical difference was observed in microglial count between Ctr and Tg mice; however, by 20 weeks, this difference was significant in both cortex and hippocampus (Fig. 1H). Lonafarnib-treated 20-week old Tg mice (Fig. 2I-L) had a significant reduction in hippocampal microgliosis (Fig. 2M) compared to age matched Unt transgenic mice.

Cortical astrogliosis was observed in transgenic mice as increased GFAP immunoreactivity (Fig. 1A; red), apparent from ~12 weeks and preceding the appearance of MC1+ neurons (supplemental Fig. 1). As transgenic mice aged, anti-GFAP immunoreactivity increased significantly in cortex, but not in hippocampus (Fig. 1I-K). Lonafarnib treatment did not consistently affect astrocytic labeling (Fig. 2N-R) possibly due to its early onset that preceded the onset of microgliosis and tau inclusions.

Next we examined the effects of lonafarnib treatment upon sumoylation and ubiquitination in rTg4510 mice. These protein modifications are prominent features of the pathology (34,36). Lonafarnib administration significantly altered the number of cells with detectable sumoylation (Fig. 2S; p=4.67×10^−5^) and ubiquitination (Fig. 2U; p=2.49×10^−3^). Using the number of double and single labeled cells, Bayesian probabilities were computed for cells labeled by MC1 given that they were labeled with anti-sumo (Fig. 2T) or anti-ubiquitin (Fig. 2V), thus comparing how well tau pathology correlated with the presence of sumoylation or ubiquitination. rTg4510 mice have many cells positive for sumoylation or ubiquitination without MC1 immunoreactivity. Using MC1 immunoreactivity as a prior, most cells also labeled positive for sumoylation or ubiquitination (supplemental Fig. 2). Treatments with lonafarnib significantly altered the Bayesian probabilities observed of MC1^+^ cells given sumoylation (Fig. 2T; p=2.71×10^−5^) and ubiquitination (Fig. 2V; p=0.016).

### Attenuation of the behavioral abnormalities in rTg4510 mice by a farnesyl transferase inhibitor

As transgenic mice age, they display progressive behavioral worsening that begins with impaired marble burial and nest shredding (26, 27) and by 30 weeks progresses to hyperexcitability, restlessness in response to human approach and obsessive circling. Using the same lonafarnib treatment protocol described elsewhere (25, 28) beginning at ten weeks of age we sought to reduce or prevent the behavioral deficits induced by tau pathology. Nest building was assessed at 20 weeks. Wild-type littermates produced well-rounded nests (Fig. 3A); however untreated transgenic mice (Tg) displayed poor nest shredding, and in some instances left bedding material completely undisturbed (Fig. 3B). Lonafarnib treatment rescued nest building (Fig. 3C, D). At five weeks, littermate controls and transgenic mice produced nests with similar scores (Ctr 3.58±0.13; Tg 3.23±0.12, p=0.639). While control mice nesting scores did not significantly change with time (p=0.840), transgenic mice scores declined (Fig. 3D). On the other hand, the marble burial deficit was observable at five weeks in transgenic mice and it remained unaltered by age (Fig. 3E-G). In contrast to nest building, the marble burial deficit (Fig. 3E, F) was not improved by the drug in the transgenic mice (Fig. 3H). One possibility that may account for this difference was the very early emergence of the marble burial deficit before lonafarnib administration, whereas shredding behavior emerged later. Lonafarnib does not appear to reverse existing pathology or deficits but seems to act as a preventive measure.

**Figure 3.**
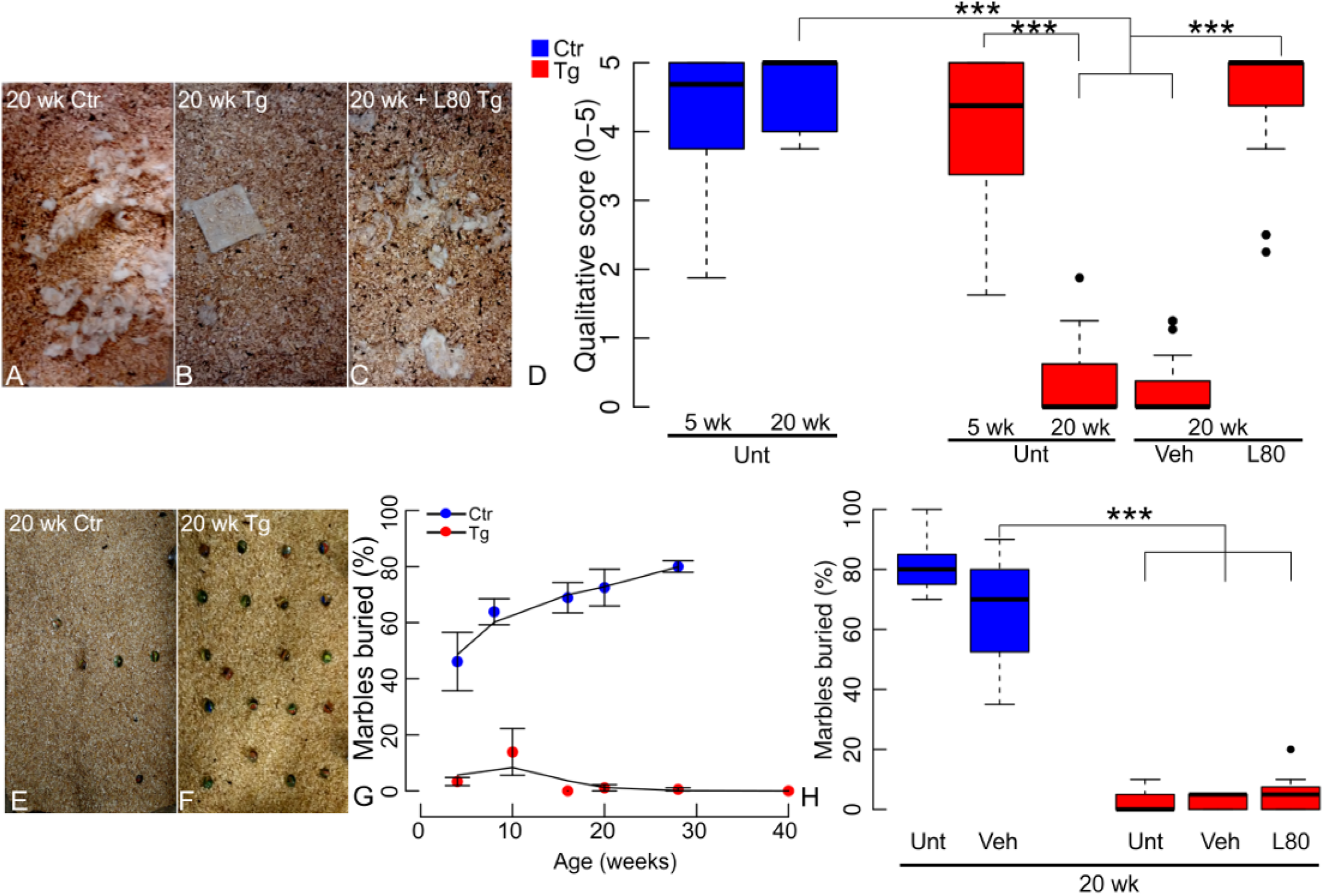
Chronic lonafarnib treatment prevents nest-building deficit in rTg4510 mice. (A) At 20 weeks littermate control mice (Ctr) display normal nest-shredding behavior, but (B) transgenic animals (Tg) failed to demonstrate nest shredding. (C) Lonafarnib treated 20-week old Tg animals shredded their nest. (D) 20-weeks old Tg mice nest quality scores averaged 0.27±0.09 in untreated mice or 0.22±0.05 in mice treated with vehicle alone whereas age-matched WT nest shredding score averaged 3.70±0.09 (both, p<1×10^−4^). Chronic and intermittent oral administration of 80 mg/Kg/day lonafarnib (L80) rescued nest building, with nest scores averaging 3.53±0.15 on treated mice. Qualitative scores were blindly assigned by observers using a scale from zero for untouched nesting material, to five for a fully shredded nest (N=6). (E) 20 week-old littermate control (Ctr) mice buried 80±2.0% of the 20 marbles in 30 min, but (F) 20 week-old transgenic (Tg) mice completely lacked digging behavior. (G) Control mice (Ctr) increased the percentage of marbles buried with age. They buried 40% of the marbles at five weeks and peaked at 30 weeks with an average of 80% of the marbles buried. Transgenic mice (Tg) fail to bury marbles as early as five weeks (two-way ANOVA genotype p<2×10^−16^, age p=0.140 n.s., interaction p=1.93×10^−5^). (H) Marble burial deficits were neither rescued by lonafarnib (L80) nor vehicle (Veh). Data is presented as boxplots of marble buried percentage per treatment group. Statistics shown for post-hoc Tukey-HSD test at ***p<1×10^−3^.

### Farnesyl transferase inhibition with lonafarnib enhanced lysosomal protein degradation

To determine the mechanism of action of lonafarnib we focused on its previously suggested role in autophagy (29). Using NIH3T3 mouse fibroblasts expressing the tandem reporter mCherry-GFP-LC3B (30) to monitor macroautophagy (supplemental Fig. 3), lonafarnib treatment resulted in a dose-dependent increase in macroautophagy flux, as revealed by an overall increase in autophagic vacuoles (AV) mostly due to an increase in autolysosome (AUT) abundance. At lower concentrations, the number of autophagosomes (APG) remained unchanged, suggesting their accelerated clearance by lysosomes (Fig. 4A-C and supplemental Fig. 3A, B). Similar results were reproduced in neuroblastoma N2a cells (supplemental Fig. 3C). The stimulatory effect of lonafarnib acted preferentially on basal macroautophagy as inferred by the fact that addition of lonafarnib to cells in which macroautophagy was induced either by paraquat (PQ) or thapsigargin (TG) did not further increase macroautophagy flux (supplemental Fig. 3D). To determine whether lonafarnib also stimulated other forms of autophagy and confirm that the observed increase in macroautophagy was not a consequence of a blockage in another degradation pathway, we used a photoswitchable reporter for chaperone-mediated autophagy (CMA) (KFERQ-PS-Dendra) (31) (Fig. 4D; supplemental Fig. 3E, F), and KFERQ-split-Venus double reporter for endosomal autophagy (8, 32) (Fig. 4E). Lonafarnib stimulated both forms of selective autophagy with maximal effect in the lower concentration range. In the case of CMA, the stimulatory effect of lonafarnib was more pronounced on basal CMA, but a significant increase was still observed after inducing CMA with PQ or TG (supplemental Fig. 3E, F). Inhibition of lysosomal proteolysis with NH_4_Cl and leupeptin in cells expressing the KFERQ-split-Venus reporter demonstrated that lonafarnib not only promoted more efficient delivery of substrate to late endosome/multivesicular bodies via endosomal microautophagy, but also stimulated efficient internalization and degradation in this compartment (Fig. 4E). Importantly, lonafarnib, had no measurable effect on proteasome-dependent degradation monitored in the same cell type using a Degron-GFP reporter (33) (Fig. 4F).

**Figure 4.**
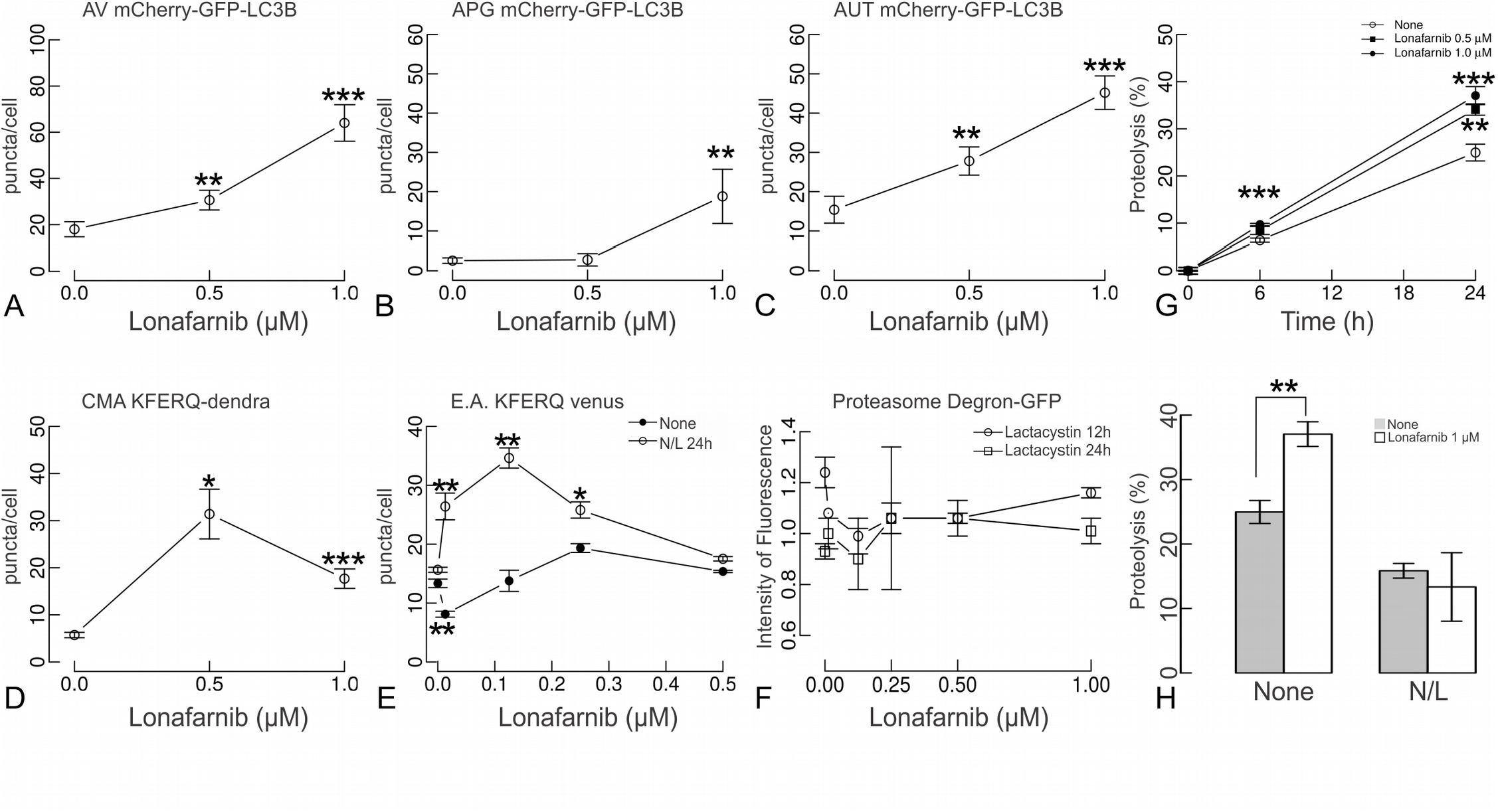
Farnesyl transferase activity inhibition activates autophagy. (A-C) NIH3T3 cells expressing the tandem reporter mCherry-GFP-LC3B were exposed to the indicated concentrations of lonafarnib for 48 h. Quantified number of autophagic vacuoles (AV), autophagosomes (APG) and autolysosomes (AUT) is shown. (D) NIH3T3 cells expressing the KFERQ-Dendra reporter were photoswitched and treated with lonafarnib as indicated above. Chaperone-mediated autophagy was quantified by the number of fluorescent puncta positive for the photoconverted Dendra per cell. (E) NIH3T3 cells expressing N and C terminus KFERQ-split Venus and treated with lonafarnib for 48 h were either treated with 20 mM NH_4_Cl and 100 μM Leupeptin (N/L) or leave untreated for the last 24 h to quantify the effect of lonafarnib on targeting (None) and degradation (N/L) by endosomal microautophagy. Quantifications in A-G were done in at least 2,500 cells per condition in three different experiments using high content microscopy. Differences with untreated (None) are significant for t-test at *p <0.05, **p <0.01 and ***p<1×10^−3^ (F) NIH3T3 cells expressing Degron-GFP and treated with lonafarnib for 48 h and supplemented with 100 μM lactacystin for the last 12 or 24 h. Proteasome-dependent degradation was calculated as the increase in the intensity of fluorescence upon lactacystin addition and after discounting the increase observed in cells treated under the same conditions but expressing a non-ubiquitinable Degron-GFP mutant. Values are expressed relative to the proteasome degradation in cells not treated with Lonafarnib that were given an arbitrary value of one. (G). Total rates of intracellular protein degradation measured in NIH3T3 cells labeled with ^3^H-leucine for 48 h. Rate of proteolysis was calculated as the percentage of the initial acid precipitable radioactivity (proteins) transformed into acid soluble radioactivity (amino acids and small peptides) at the indicated times. (H) Contribution of lysosomes to total protein degradation was analyzed supplementing cells with NH_4_Cl and leupeptin (N/L). Data is presented as mean ± s.e.m (N=6 wells in 3 independent experiments)

The overall effect in all types of autophagy suggested a direct effect of lonafarnib in endo/lysosomal compartments shared by all these autophagic pathways without influencing the proteasome. To assess this possibility, proteolysis of long-half-life proteins was evaluated after a long pulse (48 h) with ^3^H-leucine. Indeed, lonafarnib increased proteolysis in a dose dependent manner (Fig. 4G), an effect that was abolished in the presence of NH_4_Cl and leupeptin (Fig. 4H). These data support that lonafarnib treatment results in an overall improvement of lysosomal function and the pathways that mediate delivery of cargo to this compartment.

### Rhes inhibition can reduce Tau-related pathology

We sought the relevant targets of farnesyl-transferase inhibition that could account for the ameliorative effects of lonafarnib on tau pathology. Ras family members are among the prominent substrates of farnesyl transferase and this modification is required for their correct localization at the inner surface of the plasma membrane and for their biological activity. Of particular interest is the Ras family member, Rhes, because it is prenylated by farnesyl transferase (12), activates autophagy, and can modulate the aggregation state of mutant Htt by promoting its sumoylation, thereby reducing cell survival (16). To demonstrate that the effect of lonafarnib on tau pathology in rTg4510 mice could in part be accounted for by the inhibition of Rhes farnesylation, we modulated Rhes levels directly. AAV vectors (overexpression or silencing) were injected into the right amygdala of 10 week-old rTg4510 mice. When analyzed at 20 weeks (Fig. 5A-F), Rhes silencing (Rhes-miR) markedly reduced the number of MC1^+^ neurons (Fig. 5H; ANOVA cortex p=1.87×10^−4^, hippocampus p=0.002), reduced microgliosis (Fig. 5I; ANOVA cortex p=9.51×10^−5^, hippocampus p=6.75×10^−4^) and increased the coronal section area (Fig. 5G; ANOVA p=3.56×10^−4^). Therefore, Rhes inhibition recapitulated the effects of lonafarnib. Rhes over-expression did not appear to worsen the pathology (Fig. 5G-J).

**Figure 5.**
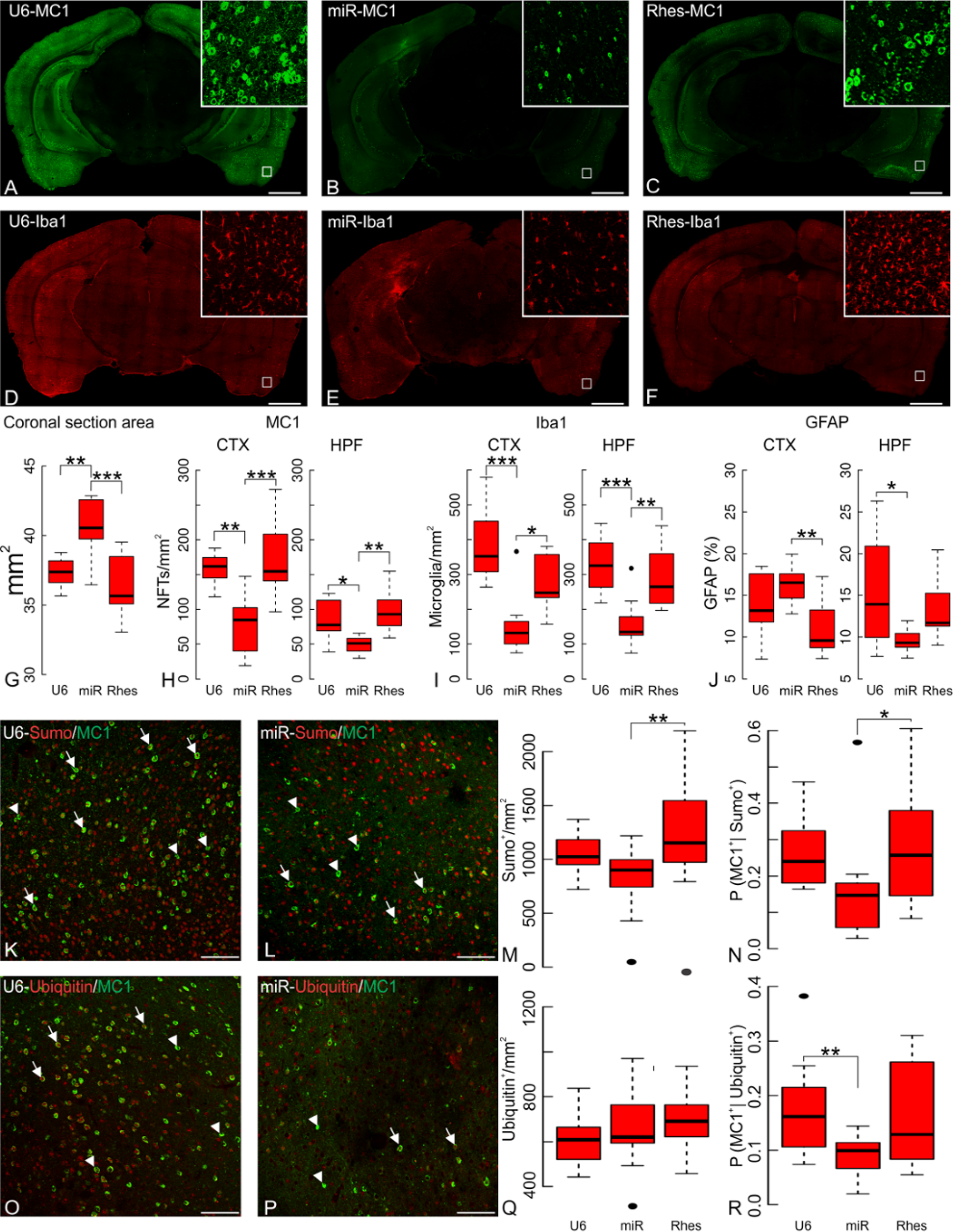
Rhes suppression mimics the protective effect of FTI in rTg410 mice. Ten-week old transgenic rTg4510 mice were intracranially infected with AAV 2/5 viral vectors containing either U6, Rhes-miR or Rhes-WT constructs and sacrificed at 20 weeks of age. (A-C) Full mosaic immunohistochemistry of MC1 immunoreactivity following Rhes suppression by Rhes-miR have fewer NFTs than transduction with U6 or Rhes. (D-F) Full-mosaic immunohistochemistry of Iba1 was also reduced in mice treated with Rhes-miR AAV. Scale bar 1 mm. Quantification of micrographs (N=3), indicates (G) significantly increased coronal section area, Rhes-miR (40.53±0.70 mm^2^) compared to U6 (37.33±0.36 mm^2^; p=5.0×10^−3^) or Rhes injected transgenic mice (36.41±0.79 mm^2^; p=3.98×10^−4^). (H) reduced number of MC1^+^ cells/mm^2^ both in cerebral cortex (CTX) and hippocampal formation (HPF). The cortical density of MC1^+^ cells/mm^2^ in animals injected with Rhes-miR was 78.76±8.875 MC1^+^ cells/mm^2^ compared to U6 (157.81±12.56 MC1^+^ cells/mm^2^; p=1.7×10^−3^) or Rhes overexpression (172.15±13.120 MC1+ cells/mm^2^; p=2.86×10^−4^). In the HPF, Rhes-miR injected mice had 49.15±7.01 MC1+ cells/mm^2^ compared to U6 (86.09±9.28 MC1+ cells/mm^2^; p=0.013) or Rhes overexpression (92.86±9.64 MC1^+^ cells/mm^2^; p=3.3×10^−3^). (I) Reduced number of microglia/mm2 in CTX and HPF in Rhes-miR injected mice. Rhes-miR injected transgenic mice were significantly reduced in the CTX (U6 control p= 5.85×10^−5^, Rhes overexpression, p=0.031) and in the HPF (U6 control p=9.48×10^−4^, Rhes over-expression, p=5.0×10^−3^). (J) Astrocytic immunoreactivity: In CTX, Rhes-miR significantly reduced GFAP immunoreactivity compared to Rhes overexpression (p=6.9×10^−3^), but not from U6 controls (p=0.357). In HPF, Rhes-miR significantly reduced GFAP immunostaining compared to U6 controls (p=0.024), but not when compared to Rhes overexpression (p=0.248); (K-M) Double immunostaining of CTX using MC1 and anti-sumo, or (O-Q) anti-ubiquitin in 20-week old rTg4510 mice and immunostained cell densities quantified as sumo^+^/mm^2^ or ubiquitin^+^/mm^2^. (M) Rhes-miR reduced sumoylated cells compared to Rhes overexpression (p=5.4×10^−3^). (Q) Rhes-miR injected mice did not show a significant reduction in ubiquitin labeled cells compared to either Rhes overexpression or U6 injected mice (p=0.615 or p=0.889, respectively). Counts of double-labeled cells in the micrographs were used to compute Bayesian probabilities of any cell to be tau immunoreactive (MC1^+^) given sumo^+^ (N) or ubiquitin^+^ (R) immunoreactivity. Rhes-miR treatments decreased these probabilities of double-labeling: (N) Rhes-miR (0.159±0.045 p=4.7×10^−3^) significantly reduced the likelihood of sumoylated cells to double-label compared to Rhes overexpression. (R) Rhes-miR (0.0907±0.0112; p=2.5×10^−3^) significantly reduced the likelihood of ubiquitinated cells to co-label compared to mice injected with U6. Statistics shown for post-hoc Tukey-HSD test at *p<0.05, **p<0.01, ***p<1×10^−3^. Scale bar 100 μm.

We examined the effects of Rhes silencing sumoylation and ubiquitination (34) in rTg4510 mice. Previous reports indicate that Rhes sumoylates mHtt aggregates (35) and tau is subject to both sumoylation and ubiquitination (34, 36). Rhes-miR significantly altered the fraction of cells positive for sumoylation (Fig. 5K-M; ANOVA p=7.73×10^−3^) and but not for ubiquitination (Fig. 5O-Q; ANOVA p=0.186). Using the number of double and single labeled cells, Bayesian probabilities were computed for cells labeled by MC1 given that they were labeled with anti-sumo (Fig. 5N) or anti-ubiquitin (Fig. 5R), thus comparing how well tau pathology correlated with the presence of sumoylation or ubiquitination. Treatment with Rhes-miR significantly reduced the double labeling with MC1^+^ of cells positive for sumoylation (Fig. 5N; ANOVA p=0.030) or ubiquitination (Fig. 5R; ANOVA p=2.7×10^−3^).

### Linking lonafarnib treatment to Rhes-mediated effects on tau

To link Rhes-induced tau pathology to its farnesylation we cultured hippocampal primary mouse neurons for three weeks at which time Rhes was over-expressed. As expected, Rhes over-expression markedly increased PHF-1 immunoreactivity. Mouse primary cultured hippocampal neurons were transduced with AAV (serotype 2/5) expressing Rhes, a GTPase inactive Rhes mutant (Rhes N33S), or a miRNA to silence Rhes (Rhes-miR). Over-expression of both Rhes variants increased PHF-1 (Fig. 6A, B). Because the Rhes GTPase mutant affected PHF-1 tau equally well as the wild type we turned our focus to Rhes as a substrate for prenylation by farnesyl transferase (37) which appears to affect its cytotoxicity in HD models (16). The farnesyl transferase inhibitor, lonafarnib added to the culture media, prevented PHF-1 tau accumulation in the presence of Rhes over-expression in a dose dependent manner (Fig. 6C, D).

**Figure 6.**
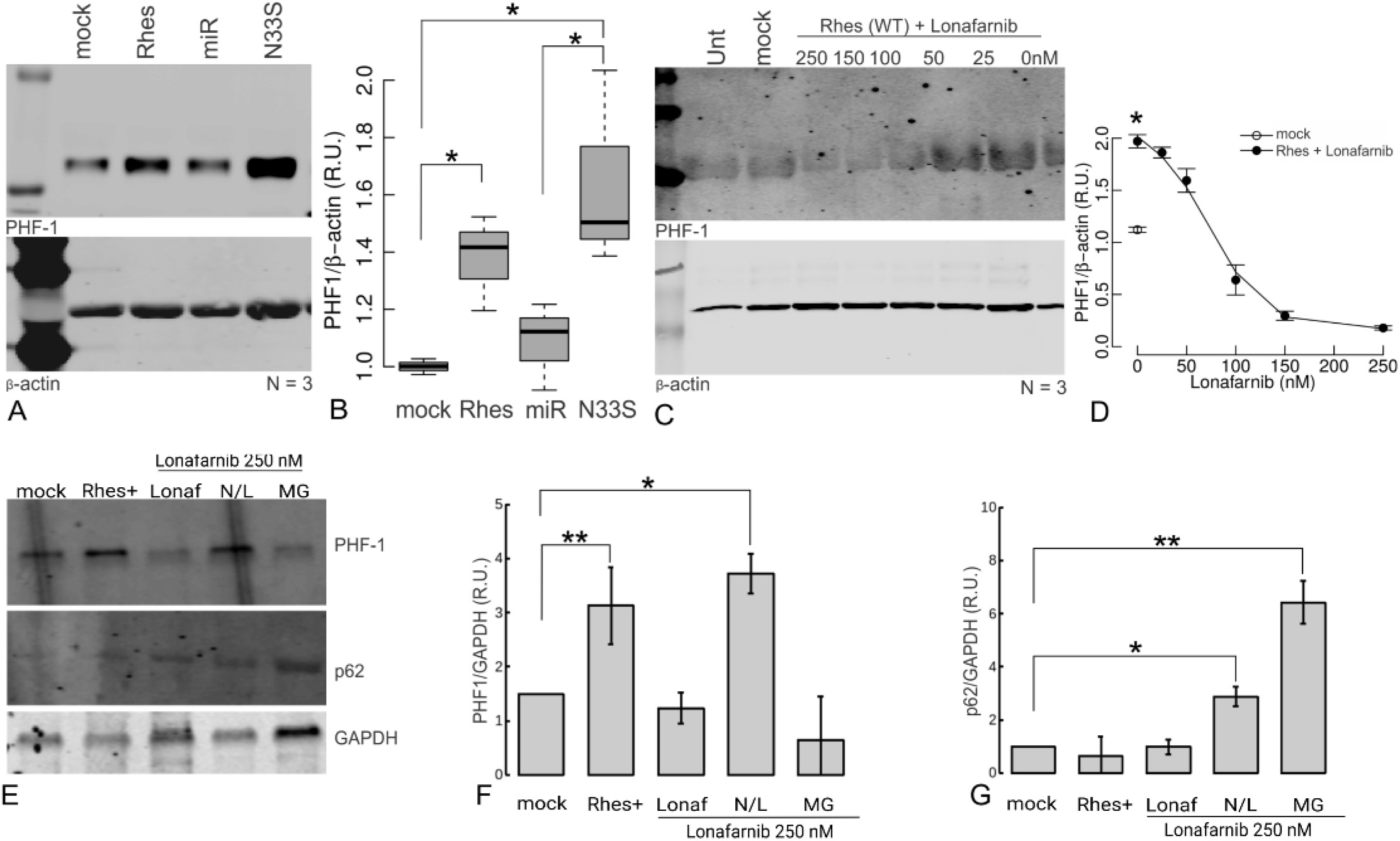
Farnesyl transferase inhibition prevents Rhes-mediated tau accumulation and activates autophagy. (A) Over-expression of either Rhes WT or the GTPase inactive mutant Rhes S33N but not Rhes silencing (Rhes miR) in hippocampal primary mouse neurons increases PHF-1 phosphorylated tau levels. A representative blot, and (B) densitometry quantification of independent replicates (N=3) is shown. (C) Farnesyl transferase inhibition with lonafarnib rescues the Rhes-induced increase in phosphorylated tau in a dose dependent manner. Representative blot, and (D) densitometry quantification of independent replicates (N=3). Beta-actin is used for loading and normalization control. Primary mouse neuronal cultures (mock) were transduced with AAV(2/5) to overexpress Rhes, and then either left untreated (Rhes^+^), or treated with 250 nM Lonafarnib alone (Lon), or additionally treated with either a cocktail containing 20 mM NH_4_Cl and 100 μM leupeptin (N/L) to block lysosomal mediated proteolysis, or with 5 μM MG-132 (MG) to block proteasome activity. Cell lysates were obtained at 12 h, or cell media and inhibitors were replenished at 12 h and lysates collected at 24 h, as indicated. (E) Proteins in the cell lysates were separated on SDS-PAGE electrophoresis and western blotted for PHF-1 S496/phospho Tau, and p62. Densitometry analysis was performed with imageJ gel analysis fuctions, and (F) the levels of PHF-1 normalized to GAPDH, or (G) the levels of p62 normalized to GAPDH were plotted as means ± s.e.m. Lonafarnib effectively prevents Rhes mediated PHF-1 increase at 24 h; and this effect can be reverted by lysosomal proteolysis blockage (N/L), but not by proteasome (MG) inhibition. Statistical analysis was performed with one-Way ANOVA, and post-hoc Tukey HSD tests were completed to determine pair-wise significance to untreated and untransduced cells. *, p<0.05. **, p<0.01. ***, p<0.001. (N=3).

We next sought to determine whether the profile of proteostasis stimulation observed for lonafarnib paralleled its effects on PHF-1 in the presence of Rhes over-expression. Hippocampal primary mouse neurons were cultured for three weeks, transduced to overexpress Rhes, in the presence of 250 nM lonafarnib, and then additionally treated with either NH_4_Cl/leupeptin that prevents lysosomal mediated proteolysis, or MG-132 a proteasome inhibitors. Indeed, as predicted, the proteasome inhibitor MG-132 had no effect on the reduction in the presence lonafarnib of the Rhes mediated PHF-1 accumulation, whereas lysosomal inhibition with NH_4_Cl and leupeptin was very effective in blocking lonafarnib action, as observed by the steadily accumulation of PHF1 immunoreactivity at 24 h (Fig. 6E, F). p62 changes are shown as positive control as it is known to undergo degradation both by the proteasome and in lysosomes (Fig. 6G).

### Developmental control over Rhes expression is associated with a homeostatic response to tau

Human induced pluripotent stem cell-derived (hiPSCs) neurons from frontotemporal mutation patients harboring either *MAPT* mutations (P301L (38), G55R (39), V337M (40), R406W (41)), a *C9ORF72* expansion (42) or clinically healthy age-matched controls were analyzed by RNAseq (Fig. 7A). Three genes were differentially expressed (DE) with the same sign in their logFC (log Fold Change) for every *MAPT* mutation line, but were not DE in the C9ORF72 (chromosome 9 open reading frame 72) line nor in any of the control lines. One of the three was *RASD2*, which encodes Rhes (Fig. 7B). (The other two were the serine-threonine kinase, *NEK9* (supplemental Fig. 4D); and the zinc-finger protein, *ZFP41* (supplemental Fig. 4E). As controls, the neuronal markers MAPT, which encodes Tau (Fig. 7C) and TUBB3, encoding pan-neuronal β-tubulin (Fig. 7D) were not DE among cells from patients carrying tau mutations. These DE genes were validated by digital RT-PCR quantification for their expression in total RNA samples taken from each of the hiPSC-derived neurons at five weeks using *GAPDH* as internal control (Fig. 7E-G, supplemental Fig. 4F-G). Further statistical validation was obtained by bootstrapping ANOVA tests (k=1000) of their CPM counts (43) (bootstrapped p-values: RASD2=8×10^−3^, ZPF41=0.047, NEK9=0.348 n.s.). *RASD2* levels in the tau mutants were confirmed in a set of isogenic controls. Real time RT-PCR in three independent sets of hiPSCs derived neurons with different MAPT mutations and their isogenic controls showed the expected DE *RASD2* levels (MAPT-P301L/P301S (Fig. 7H), *MAPT-V337M* (Fig. 7I) and *MAPT-*R406W (Fig. 7J). In agreement with the RNAseq, *MAPT* expression did not change significantly in any of isogenic controls (Fig. 7K-M). Curiously, however, the direction of change in *RASD2* was decreased in the tau mutants relative to the controls.

**Figure 7.**
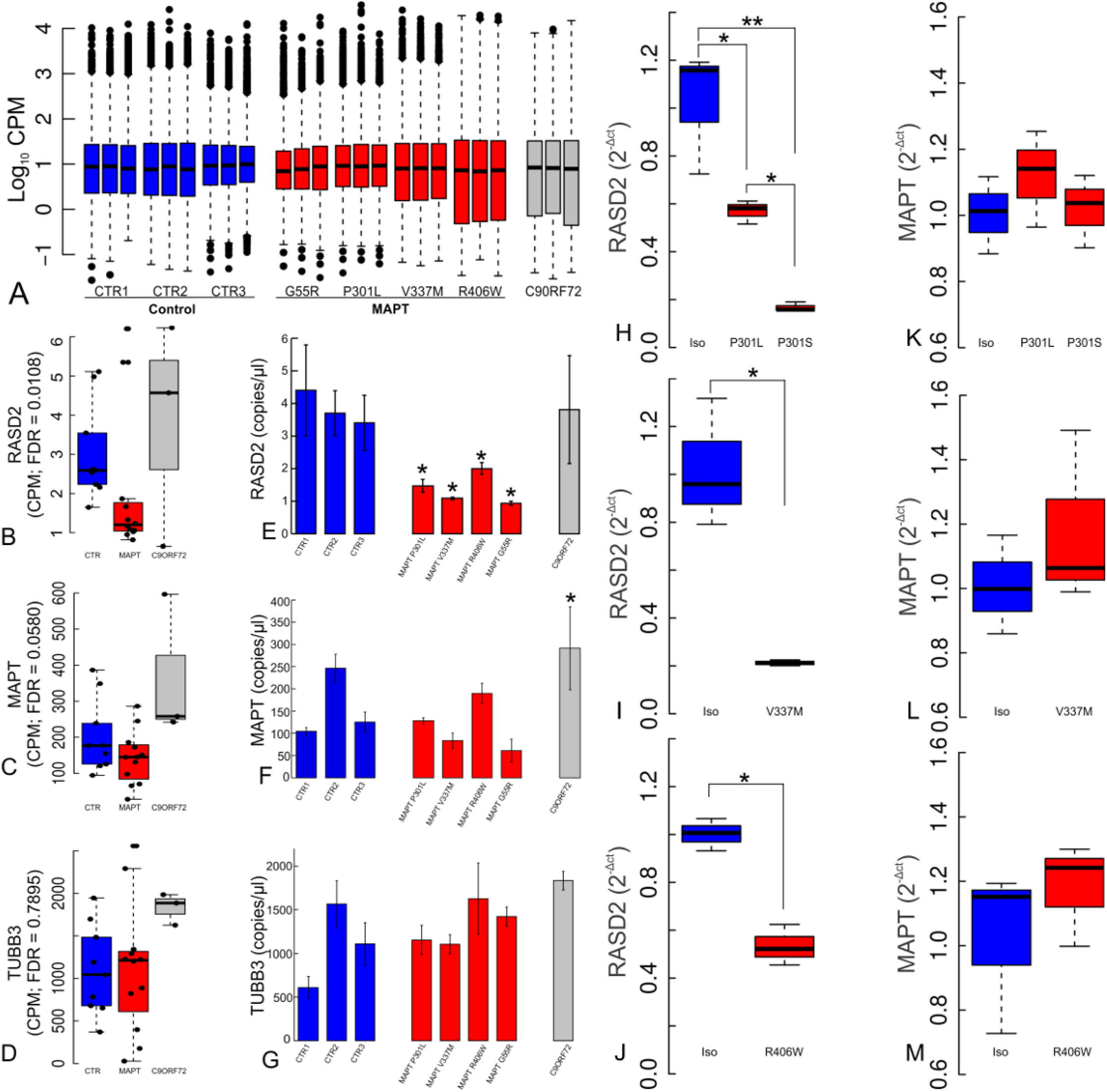
RASD2 is reduced in transcriptome profiles of hiPSCs-derived neurons from FTD patients carrying MAPT mutations. (A) RNAseq hiPSCs-derived neurons from FTD patients carrying *MAPT* mutations (G55R, P301L, V337M, and R406W) or a *C9ORF72* expansion and three age-matched healthy controls in culture for five weeks. Boxplots of normalized CPMs per library are shown. (B) RASD2 was commonly deregulated across each patient line carrying *MAPT* mutations. (C) *MAPT* expression does not change across *MAPT* mutation carriers. (D) TUBB3 is highly expressed and remains unaltered across samples. Digital PCR data (E-G) presented as mean copies/μL ± s.e.m. validated the findings of each of these genes for each cell line studied. RASD2 suppression in the presence of tau mutations compared to their respective isogenic control lines was verified by Taqman RT-PCR, using (H) F0510 cells harboring P301L and P301S mutations (ANOVA p=1.45×10^−3^), (I) Cells harboring MAPT-V337M (t-test p=0.035), and (J) cells harboring MAPT-R406W (t-test p=2.14×10^−3^). (K-M) *MAPT* expression remained unchanged in the *MAPT* mutations and their corrected isogenic lines.

To understand the context of *RASD2* level regulation we next found that *RASD2* expression levels in a control line increased during iPSCs differentiation; however, in *MAPT-P301L* and *MAPT-V337M* lines *RASD2* levels remained low throughout differentiation (Fig. 8A). Over this same time period, *MAPT* expression increased and showed no statistical difference between control cells and the *MAPT* mutants (Fig. 8B). Thus, under control conditions *RASD2* levels rose coincident with increased tau levels, but were suppressed in the presence of tau mutations.

**Figure 8.**
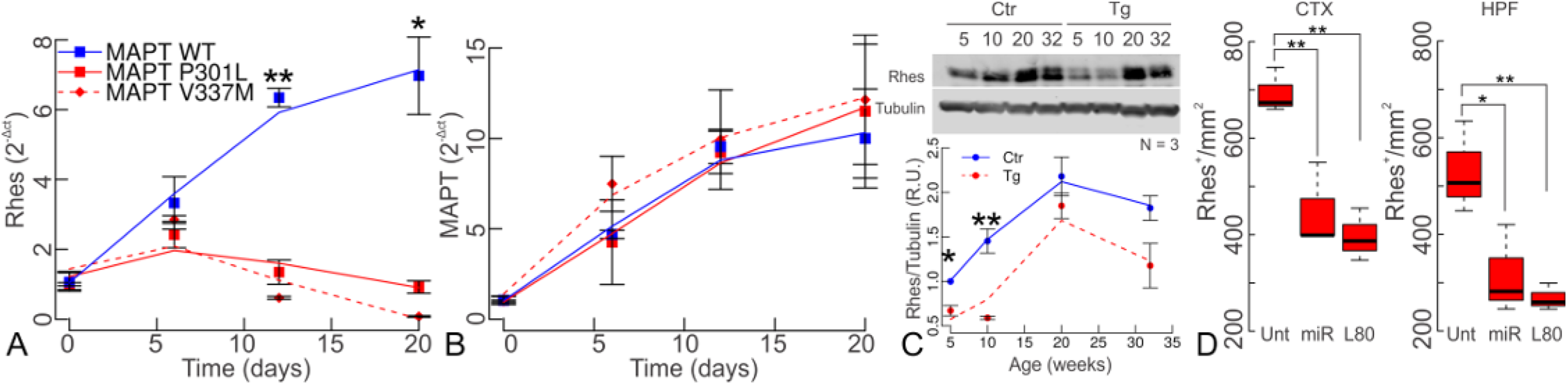
Rhes levels increase as neurons age can be prevented by Tau mutation. (A) Rhes expression is reduced in *MAPT-P301L* and MAPT-V337M derived neurons during hiPSC neuronal differentiation as early as the neurorosette stage but increases during cell differentiation in MAPT-WT cells. (B) *MAPT* levels increase continuously during neuronal differentiation regardless of *MAPT* genotype. (C) Rhes quantification in rTg4510 transgenic mice (Tg) and littermate controls (Ctr). While Rhes levels increase as mice age in both transgenic and control mice, these are significantly lower in younger transgenic mice (N=3; two-way ANOVA, age p=3.24×10^−3^; genotype p=8.01×10^−3^; interaction p=0.88 n.s.). (D) Reduction of Rhes in 20 week-old transgenic mice. Untreated rTg4510 mice had a mean of 693±27 Rhes+ cells/mm^2^ in the cortex (CTX) and 449±15 cells/mm^2^ in the hippocampus (HPF). Chronic administration of lonafarnib reduced Rhes to 449±50 cells/mm^2^ (p=8.94×10^−3^) in the CTX and to 316±51 cells/mm^2^ (p=0.035) in the HPF. Rhes-miR reduced Rhes to 396±31 cells/mm^2^ (p=3.41×10^−3^) in the cortex and to 268±16 cells/mm^2^ (p=0.015) in the hippocampus.

Because iPSC-derived neurons remain relatively immature, tracking *RASD2* levels at more advanced ages is not possible on them. Therefore, levels of the *RASD2* transcript were measured in rTg4510 transgenic mouse (22) brains and their littermate controls at 5, 10, 20 and 32 weeks. Consistent with cultured cells, as mice aged, Rhes levels increased (Fig. 8C). Rhes levels were significantly lower in younger transgenic mice (5 and 10 weeks) that expressed mutant tau than their littermate controls that expressed only wild-type mouse tau. However, by 20 and 32 weeks the difference between transgenic and littermate controls was no longer significant. These observations suggested that a compensatory Rhes reduction could protect younger animals by maintaining lower Rhes levels and this homeostatic response is overcome as pathology progresses. Lonafarnib treatment or Rhes-miR AAV reduced Rhes levels in rTg4510 mice (20 weeks), in cortex (Fig. 8D; ANOVA p=3.0×10^−3^) and hippocampus (ANOVA p=0.014).

## Discussion

Attempts to treat the tauopathies, particularly those individuals with tau mutations, has been impeded by the absence of an understanding of the molecular pathways that connect genotype to phenotype. The remarkable effects of the farnesyl transferase inhibitor, lonafarnib, on tau pathology allowed the use of this drug as a probe for the discovery of disease-relevant pathways. Although farnesyl transferase inhibitors, such as lonafarnib, will affect many farnesylated substrates, the major effects of the drug appear to be mediated through membrane associated Rhes and therefore points to a regulated pathway as a druggable target. Lonafarnib has its principal effect on enhancing protein degradation in the lysosome (Fig. 4). When Rhes was over-expressed in neurons to increase PHF-1 and then treated with lonafarnib to reduce PHF-1, the effects of the drug were prevented by the lysosomal inhibitors, NH_4_Cl and leupeptin (Fig. 6E-G). Furthermore, inhibition of Rhes in the rTg4510 mouse phenocopied the effects of lonafarnib (Fig. 5). Positing the lysosome as the drug target in the mouse is consistent with observations that the P301L mutation interferes with the degradation of tau through all the autophagic pathways (8).

Many questions remain regarding the pathway from membrane disassociation of Rhes to lysosomal activation (supplemental Fig. 5). Farnesylation of Rhes may serve as a switch between its effect on mTOR to inhibit autophagy and on beclin-Bcl2 to activate autophagy. The regulation of autophagy by Rhes under neurodegenerative disease conditions was reported to be independent of mTOR (15), and as such, strategies that activate mTOR may not be opportune therapeutic options. On the other hand, strategies that activate autophagy have shown beneficial effects (44, 45). Contributing to the mechanism of Rhes is its role as a sumo ligase (16). Tau is both sumoylated (34) and ubiquitinated (36, 46). Although a direct interaction between Rhes and tau has not yet been demonstrated, mutant Huntingtin can bind to Rhes (16).

Sumoylation can become activated under certain stress conditions. An increase in Sumo conjugation has been observed, for example, upon heat shock, osmotic stress and hibernation (47–49). Both Rhes-miR and lonafarnib were able to suppress sumoylation and ubiquitination in rTg4510 mice (Fig. 2S-V, and Fig. 5K-R). The presence of many more sumo^+^ than MC1^+^ cells suggests that a potentially important facet of the pathology may occur in cells undergoing sumoylation or ubiquitination without tau inclusions, and that activation of these pathways is conducive to the induction of neurofibrillary pathology. Both lonafarnib (Fig. 2T, V) and Rhes-miR (Fig. 5N, R) decreased the probabilities of double-labeling, thereby suggesting a link between the Rhes pathway and activation of sumoylation and/or ubiquitination.

From stem cells (50) to aged cells (51) proteostasis is a highly regulated process over the lifespan. Indeed, Rhes levels are developmentally regulated with low levels in neuronal precursors and young animals that gradually increase with maturation (Fig. 8). The persistently low levels of Rhes in tau mutation harboring iPSC-derived neurons (Fig. 7) suggested a homeostatic response to the mutation (15, 52) by which the cell can sense tau mutations before inclusions are evident and respond by activating the lysosome through endogenous reduction of Rhes. In fact, an increase in lysosomal proteolysis in response to mutant tau expression has been reported (8). Immature hiPSC-derived neurons and young rTg4510 mice appear to compensate for the deleterious effects of the tau mutation by reducing Rhes levels (Figs. 7B, E, H-J, and 8C). Under control conditions *RASD2* (the Rhes transcript) expression rises coincident with increases in tau levels; whereas, this rise in Rhes fails to occur in the presence of tau mutations (Fig. 8A-B). Tracking Rhes levels in hiPSC-derived neurons is limited by the fact that these cells do not fully mature. However, the advancing tau pathology in rTg4510 mice, appeared to escape auto-regulatory controls and Rhes levels do rise (Fig. 8C). This Rhes sensor system appears sufficiently sensitive to detect the tau P301L mutation in the alternatively spliced tau exon 10, which is expressed, albeit at low protein and mRNA levels in hiPSC-derived neurons (unpublished observations). A similarly complex picture regarding the role of Rhes in HD has been proposed (15). Rhes is down-regulated in HD (53, 54) and yet increasing Rhes expression augments cytotoxicity (38). Thus the Rhes pathway serves as a highly sensitive sensor. This Rhes lowering strategy was adopted pharmacologically in older animals to treat the tauopathy. By inhibiting Rhes farnesylation its levels are reduced probably due to its degradation when free in the cytoplasm.

The pharmacological approach described here will likely require early and chronic intervention. Indeed, once pathology emerged lonafarnib was no longer effective. Impairment in marble burial, which emerged before 10 weeks when the drug was delivered and well-before tau inclusions were observed could not be remediated. Nevertheless, the broad implication of proteostasis in many neurodegenerative diseases including Parkinson’s disease, Huntington’s disease, Alzheimer’s disease, and other types of frontotemporal dementia (51, 55, 56) requires a concerted effort to explore relevant pathways for therapeutic effects. The safety profile of lonafarnib (19, 20) and its efficient crossing of the blood brain barrier (19), makes this drug a promising candidate for re-purposing to treat tauopathies due to tau mutations.

## Materials and methods

### Animal maintenance and treatments

This study was conducted in accordance with the National Institute of Health Guide for the Care and Use of Laboratory Animals, the Association and Accreditation of Laboratory Animal Care International (AAALAC) guidelines, and under the authorization of the Institutional Animal Care and Use Committee and the University of California Santa Barbara. Animals were placed on a standard rodent diet ad libitum, and housed under a 12 h/12 h light-dark cycle.

Transgenic rTg4510 mice (22) were purchased from Jackson Laboratory (Bar Harbor, ME) and maintained by breeding B6 *CaMKII-tTA* heterozygotes (003010) with FVB.tetO-MAPT-P301L heterozygotes (015815). Weanlings were PCR-genotyped with DNA obtained from an oral swab (Puritan, Guilford, ME) using Kapa mouse genotyping kits for DNA extraction and amplification (Kapa Biosystems, Wilmington, MA) as suggested by the Jackson Laboratory (PCR primers used, forward: oIMR8746, tTA reverse: oIMR8747, tTA control forward: oIMR8744, tTA control reverse: oIMR874, *MAPT* forward: 14159, *MAPT* reverse: 14160, *MAPT* control forward: oIMR7338, *MAPT* control reverse: oIMR7339). Identified double transgenics were used in the experiments and reported here as transgenics (Tg); littermates identified as negative for both tTA and *MAPT-P301L* were used as non-transgenic controls (CTR) where indicated. Doxycycline was not used to turn off the expression of MAPT-P301L transgene in this study.

Lonafarnib was synthesized by Cayman Chemical Company (Ann Harbor, MI) and certified 99.9% or higher purity by HPLC, and stored lyophilized at −20°C. 100X lonafarnib was dissolved in DMSO heated to 95°C until solution was clear and 12 mg/mL was re-suspended in 20% (2- Hydroxypropyl)-β-cyclodextrin (HBCD; Sigma-Aldrich) as a vehicle (Veh). Re-suspended solution was stored at 4°C for less than a week. 10 week-old rTg4510 mice were orally administered a dose of 80 mg/Kg/day of lonafarnib, or vehicle alone by gavage-feeding in an alternate schedule of five consecutive days followed by five resting days for 10-weeks.

Overexpression and silencing of Rhes was achieved by sterotactic injections of ice-cold lactated ringer dialyzed adeno-associated viral vectors (AAV 2/5) carrying either Rhes-IRES-GFP (Rhes), or an engineered Rhes-targeted synthetic microRNA (Rhes-miRNA) or U6 alone, to the amygdala on bregma coordinates −0.30 (AP), −2.00 (L) and −4.80 (DV). Stereotactic injections of 1×10^9^ viral particles were performed on rTg4510 mice at ten weeks of age. Packaged viral vectors were kindly provided by Beverly Davidson, University of Pennsylvania and kept at −80°C until dialysis.

### Sample Preparation and Immunocytochemistry

Animals were transcardially perfused using 4% paraformaldehyde in 0.1 M sodium cacodylate (Electron Microscopy Sciences, Hatfield, PA) for 15 min at room temperature. Brains were then dissected and immersion fixed for 48 h at 4°C. Immunocytochemistry was carried as described elsewhere (63). Briefly, samples were rinsed 5 x 5 min in phosphate buffered saline (PBS; pH 7.4) and then coronally sectioned at 100 μm using a vibratome (Leica, Lumberton, NJ). Sections were immersed in normal donkey serum 1:20 in PBS containing 0.5% BSA, 0.1% Triton-X 100, and 0.1% sodium azide (PBTA) at 4°C on a rotator for continuous overnight agitation followed by immersion MC1 (provided by Peter Davies, Feinstein Institute for Medical Research, Manhasset, NY; mouse monoclonal, 1:200), anti-Iba1 (Wako laboratory chemicals, Richmond, VA; rabbit polyclonal, 1:200), anti-GFAP (abcam, San Francisco, CA; chicken polyclonal, 1:500), anti-ubiquitin (abcam, ab7780; rabbit polyclonal), and anti-sumo 1 (abcam, ab11672; rabbit monoclonal) diluted in PBTA. The following day sections were rinsed in PBTA 5 x 5 min, 1 x 1 h in PBTA and then placed in secondary antibodies donkey anti-mouse 568, anti-rabbit 488, and anti-chicken 647 (Jackson ImmunoResearch Laboratories; West Grove, PA; 1:200). Finally, secondary antibodies were rinsed and mounted using Vectashield (Vector Laboratories Inc., Burlingame, CA) on a glass slide and sealed under an 18 x 18 #0 micro coverslip (Thomas Scientific; Swedesboro, NJ) using nail polish.

### Large-scale mosaic acquisition, registration and image analysis

Specimens were screened and imaged using an Olympus Fluoview 1000 laser scanning confocal microscope (Center Valley, PA) equipped with an argon 488 argon photodiode laser, and HeNe 543/633 photodiode lasers as well as precision motorized stage (Applied Scientific Instrumentation, Inc. Eugene, OR). Three coronal sections per mouse that spanned the hippocampal formation from anterior to posterior were selected. Mosaics were captured using an UPlanSApo 20x air lens, N.A. 0.75 at 1 μm intervals along the z-axis and a pixel array of 800 x 800 in the x-y axes. Image stacks were collected sequentially using the Olympus Flouview software version 4.2 with 5% overlap between individual tiles. Alignment and registration of individual tiles were performed in a semiautomated fashion by *Imago* 1.5 (Mayachitra Inc. Santa Barbara, CA). All digital image datasets used for spatial analyses have been deposited to Bisque (Bio-image Semantic Query User Environment) (64) and are publically available for further use with permission: https://bisque.ece.ucsb.edu/client_service/view?resource=https://bisque.ece.ucsb.edu/data_service/00-iZXFv5ELXaikLmi2viaSRJ.

Whole brain mosaics were manually segmented into either hippocampal or cortical regions using FIJI version 2.0.0. Images were set as 8-bit and thresholds were manually adjusted. Total number of MC1^+^ neurons and Iba1^+^ microglia were quantified using FIJI’s analyze particles function (65). Segmentation of anti-GFAP staining was performed using WEKA (66), a third party library included in FIJI. The number of Rhes^+^, sumo^+^ and Ubiquitin^+^ cells were quantified in FIJI by discretely counting cells positively immunostained over threshold adjustments indicated above, on confocal z-stacks of micrographs collected at 1024×1024 pixel arrays.

### Behavioral characterization

Nest shredding behavior was qualitatively measured according to published guidelines (27) with modifications. Briefly, unscented nestlets were provided during husbandry and kept in the cage for 24 h. Nest shredding behavior overnight was estimated on a 0-5 scale with zero indicating unshredded bedding material while a five denoted a completely shredded nest that displayed a rounded appearance. Twenty-five blinded independent observers reviewed images of overnight nests and the average quality scores were calculated. Following this time, a picture of the remaining nestlet or the nest was taken and the pictures scored on a 0 to 5 scale: 0 for an un-shredded nest, 1 and 2 for nests slightly to moderately shredded, 3 and 4 for nests that were shredded but the appearance of the nest was flat, and 5 for nests that were fully shredded and the appearance of the nest was round and full. Scores of each nest were assigned blindly by 25 independent observers, and these scores were averaged to produce net-scores (Fig 3D). One-way ANOVA tests followed by Tukey post-hoc test were used to determine statistical significance in the difference of means.

Marble burial was evaluated as previously reported (26) with minor modifications as follows: 20 marbles of 15 mm in diameter were spaced by 4 cm in five rows of four marbles each, on the surface of a gently packed 5 cm deep wood chip bedding in a double-size rat cage. A mouse was left alone in the cage for 30 min and then returned to its housing cage. The number of marbles buried was counted by an observer blinded to the treatment. Any marble buried more than 2/3 of its size was counted. Each mouse was assessed three times on consecutive days, at the same time in the afternoon, and data reported as average ± s.e.m. of the percentage of buried marbles per animal. One-way or two-way ANOVA tests followed by Tukey post-hoc where applicable were used to determine statistical significance.

### Evaluation of intracellular proteolysis and autophagy in lonafarnib treated cells

Mouse fibroblasts (NIH3T3) or neuroblastoma N2a cells were obtained from the American Type Culture Collection (ATCC). Cells were maintained in Dulbecco’s modified Eagle’s medium (DMEM) (Sigma, St. Louis, MO) in the presence of 10% newborn calf serum (NCS), 50 μg/ml penicillin and 50 μg/ml streptomycin at 37°C with 5% CO2, and treated with varying concentrations of lonafarnib (0.25-1 μM).

Macroautophagy activity in intact cells was measured upon transduction with lentivirus carrying the mCherry-GFP-LC3 tandem construct (30). Cells were plated on coverslips or glass-bottom 96- well plates and fluorescence was read in both channels. Puncta positive for both fluorophores correspond to autophagosomes whereas those only positive for the red fluorophore correspond to autolysosomes. Autophagic flux was determined as the conversion of autophagosomes (yellow) to autolysosomes (puncta; red).

Chaperone-mediated autophagy (CMA) activity was quantified in intact cells using the photoswtichable (PS) KFERQ-PS-Dendra2 reporter transduced into cells using lentiviral delivery (31). Cells were photoswitched with a 405 nm emitting diode (LED: Norlux) for 4 min with the intensity of 3.5 mA (current constant). CMA activity was quantified as number of Dendra photoconverted positive puncta per cell.

Endosomal microautophagy (32) activity was measured in intact cells using a recently developed KFERQ-N-terminus Split Venus and the KFERQ-C-terminus split Venus (8). As the half proteins are targeted to the multivesicular bodies in late endosomes, the confined space inside these vesicles favors the interaction of the two parts, and once Venus is joined punctate fluorescence results. The amount of reporter undergoing degradation in this compartment can be estimated by comparing number of puncta in cells treated or not with NH_4_Cl and leupeptine to inhibit lumenal degradation.

To determine possible changes in proteasome-dependent degradation, cells were transduced with lentivirus carrying the degron-GFP (33) that will undergo rapid degradation inside cells unless lactacystin is added to inhibit proteasome activity. Rates of degradation are calculated as the increase in total cellular fluorescence upon addition of lactacystin, and discounting changes in cells transduced with lentivirus carrying a mutant-degron-GFP, unable to undergo selective proteasome degradation.

For all reporters, cells plated in glass-bottom 96-well plates were treated for the indicated times and after fixation images were acquired using a high-content microscope (Operetta, Perkin Elmer). Images of 9 different fields per well were captured, resulting in an average of 2,500-3,000 cells. Nuclei and puncta were identified using the manufacturer’s software. The number of particles/puncta per cell was quantified using the “particle identifier” function in the cytosolic region after thresholding in non-saturated images (67). In all cases, focal plane thickness was set at 0.17 μm and sections with maximal nucleus diameter were selected for quantification. Values are presented as number of puncta per cell section that in our acquisition conditions represents 10-20% of the total puncta per cell.

To measure degradation of long-lived proteins, confluent cells were labeled with ^3^H-leucine (2 μCi/ml) for 48 h at 37°C, and then extensively washed and maintained in complete (10% NCS) or serum-deprived media containing an excess of unlabeled leucine (2.8 mM) to prevent reutilization of radiolabeled leucine (68). Aliquots of the media taken at different times were precipitated with TCA and proteolysis was measured as the percentage of the initial acid-insoluble radioactivity (protein) transformed into acid-soluble radioactivity (amino acids and small peptides) at the end of the incubation. Total radioactivity incorporated into cellular proteins was determined as the amount of acid-precipitable radioactivity in labeled cells immediately after washing.

All numerical results are reported as mean + s.e.m. Statistical significance of the difference between experimental groups was analyzed by two-tailed unpaired Student’s t-test. Differences were considered statistically significant for p<0.05.

### Evaluation of proteolysis contribution on lonafarnib enhanced tau degradation

Hippocampal primary mouse neuronal co-cultures were prepared from E19 mouse hippocampi, dissected and dissociated with 2.5% trypsin, plated in Poly-L-Lysine coated cell culture six-well plates at a density of 250,000 neurons per well. Hippocampal neurons were matured for three weeks in neurobasal media supplemented with N2, B-27 and L-glutamine with 1% Penicillin and Streptomycin cocktail. Neurons were fed by replacing half of the culture media with pre-warmed and freshly supplemented media twice per week. Cells were then transduced with 1×10^9^ AAV2/5 particles per well to either over-express or silence Rhes, and simultaneously treated with Lonafarnib at varying concentrations ranging from 0 to 1 μM. Cell lysates were obtained 24h post-treatment with RIPA buffer after adding proteasome inhibitor cocktail (cOmplete, Roche, Bradford, CT), and phosphatase Inhibitor coctail (Sigma-Aldrich), per the manufacturer’s instructions. Proteins were separated with SDS-PAGE, and wet-transferred to nitrocellulose. Membranes were then blotted with PHF-1 and beta-actin (1:1000, each) and imaged using Li-COR fluorescent scanner.

To dissect the contribution of diverse proteolysis pathways, hippocampal primary mouse neurons treated as described above with AAV2/5 to overexpress Rhes in the presence of 250 nM Lonafarnib were further treated in the presence of either a cocktail of 20 mM NH_4_Cl and 100 μM Leupeptin to prevent lysosomal mediated proteolysis, or 5 μM MG-132 to block proteasome mediated proteolysis, or left untreated. Cell lysates were collected as described above 12 h after treatments, or treatments were replenished at 12 h, and lysates were collected 24 h after treatments began. Membranes were blotted with PHF-1, p62 (ab56416, abcam) at 1:1000 or GAPDH (MAB374, Millipore) at 1:10000.

### hiPSCs and neuronal differentiation

All work presented here using human derived cell lines was reviewed and approved by oversight committees: Washington University in St. Louis, IRB No. 201306108, and University of California Santa Barbara, hSCRO: 2017-1065. hiPSCs (human induced pluripotent stem cells) were reprogrammed from skin biopsy fibroblasts from three independent clinically healthy controls (CTR2, CTR3, and F11349.2), and four independent fronto-temporal mutation patients carrying different MAPT mutations (P301L, G55R, V33M, or R406), and a disease control, C9ORF72 (supplemental Table 1). hiPSCs were maintained and propagated in Matrigel (BD Byosciences, San Jose, CA) coated six-well plates in mTSER supplemented (Stem Cell Technologies, Vancouver, Canada) KO-DMEM/F-12 media (ThermoFisher, Carlsbad, CA). Every hiPSC line was karyotyped as normal except control line F11349.2, which had a 3q27 translocation of unknown origin (Cell line genetics, Madison, WI). The cell lines were gender-randomized: females– control (CTR3), *MAPT* (V337M, R406W and G55R), and *C9ORF72;* males– controls (CTR2 and F11349), *MAPT* (P301L). Every hiPSC line was validated with immunohistochemistry for expression of *Sox2* (AB5603, Millipore, Temecula, CA), *NANOG* (AB5731, Millipore) and *OCT4* (ab18976, abcam, Cambridge, MA). Neurons were derived from hiPSCs by culturing colonies in the presence of 5 μM dorsomorphin and 10 μM SB431542 (Sigma-Aldrich, St. Louis, MO) in DMEMF12 media supplemented with 20% KOSR (ThermoFisher), 1X non-essential aminoacids, 1X GlutaMAX and 55 μM β-mercaptoethanol (ThermoFisher). Supplemented media was replenished on days 3 and 5, after which media is mixed 1:1 with neural-induction media (NIM: DMEMF12 supplemented with 1X N2, 1X non-essential amino acids, and 1X GlutaMAX (ThermoFisher). After day 5, SB431542 was withdrawn, but dorsomorphin supplementation was maintained to day 9, when media was changed to NIM supplemented with 0.25 μg/mL laminin from human placenta (Sigma-Aldrich) and 8 ng/ml bFGF (Peprotech, Rocky Hill, NJ). This media is replenished every other day until day 15. Neurorosettes were microdissected, grown and propagated in suspension as neurospheres in NIM supplemented with 8 ng/ml bFGF and 1X B27 media. These neurospheres were passaged weekly for up to four weeks and then enzymatically split into single cells with 1:1 accutase trypsin (ThermoFisher), and differentiated to neurons on poly-L-ornithine/laminin (Sigma-Aldrich) coated six well plates. Neurons matured for five weeks in Neurobasal media supplemented with B27, 1X GlutaMAX (ThermoFisher), and a cocktail of neurotrophic factors containing 10 ng/mL of each NT3, BDNF, and GDNF (ThermoFisher). Neurons were characterized with immunohistochemistry for expression of *MAP2* (4542; Cell Signaling Technology, Danvers, MA), tau (PHF-1; provided by Peter Davies, Feinstein Institute for Medical Research, Manhasset, NY), synapsin (AB1543, Millipore) and *PSD95* (ab13552, abcam). Antibiotics were not added to the media for the cell culture studies.

### Isogenic iPSC and neuronal differentiation

iPSCs carrying MAPT-P301L (F05010), MAPT-V337M (ND32591A), and MAPT-R406W (F11362) mutations were differentiated to neurons. Isogenic lines were generated in a footprint-free, seamless manner (i.e. no blocking mutations) using CRISPR/Cas9. Sanger sequencing was performed to confirm clones were correctly edited at on-target sites and free of indels or modifications at predicted off-target sites. All clones were confirmed to have normal karyotype after editing. Isogenic pairs were differentiated into neurons using a two-step protocol. Briefly, iPSCs were plated in neural induction media (StemCell Technologies) in a 96-well v-bottom plate to form highly uniform neural aggregates and after five days transferred onto culture plates. The resulting neural rosettes were then isolated by enzymatic selection. The NPCs are differentiated in planar culture in neural maturation medium (neurobasal medium supplemented with B27, GDNF, BDNF, cAMP) for six weeks

### mRNA Transcript quantification

Genome-wide poly-A RNA transcripts were quantified using five-week-old neuronal cultures differentiated from patient and control hiPSCs as described above. RNA sequence was obtained from three independent hiPSC neuronal differentiation events from each individual. Total RNA was collected from a total of four six-well plates of neurons grown at a density of 2×10^5^ cells/well (miRVANA kit, ThermoFisher). Poly-A yield was enriched from 5 μg of total RNA using poly(A) purist MAG kit (ThermoFisher). The RNA fraction was used to prepare libraries for RNAseq with an average size of 200 bp. (Ion Total RNA-seq Kit v2, Life technologies, Austin, TX). Libraries were quantified using Qubit dsDNA HS Assay kit (ThermoFisher) and their sizes measured in a Fragment Analyzer (Advanced Analytical, Ankeny, IA) using NGS Fragment Analysis Kit (Agilent genomics, Santa Clara, CA), 2.1 x10^−3^ pmol of cDNA per library were sequenced by ion semiconductor sequencing (Ion Torrent, ThermoFisher) per the manufacturer’s instructions obtaining an average of 3.5×10^7^ reads per library.

The average size of the sequencing reads after adapter and index clipping was 80 bp. The resulting FASTQ files were mapped to the human genome (hg19) using the tophat suite (version 2.1.0) (57). Reads mapped were filtered for quality control including removal of PCR duplicates, reads with low mapping scores, or mapping repetitive sequences, ribosomal or nucleolar sequences. Sorted BAM files were further processed with Samtools (version 1.20) (58) and Bedtools (version 2.24) to produce tabulated counts of reads aligned to the consensus coding sequences (CCDS, release 21 (59) Reads mapped to CCDS were normalized on calculated library sizes by TMM with bioconductor edgeR suite (R version 3.0.1, edgeR version 3.2.4 and limma version 3.16.7) (60–62) to obtain CPMs on a gene basis per library. Normalized reads were filtered for genes with < 5 counts in three independent libraries or fewer. All libraries obtained presented gene expression distributions that approach a log-normal distribution, as expected. Furthermore, the distribution of normalized gene expression by logarithmic CPMs was comparable across each library sequenced (Fig 1A).

Differential expression (DE) analysis was conducted with a generalized linear model (GLM) using edgeR, and fit with all control lines compared to either all MAPT carriers or to each patient independently. DE genes were selected with FDR <0.05.

Our library preparation cannot distinguish isoform specific DE unequivocally, therefore we report differences computed on full-length transcripts. Tabulations for all DE genes observed for each comparison, with their corresponding expression levels (CPM), logarithm fold change (LogFC), FDR and Benjamini corrected p-values are included in the supplemental data. Set intersections for DE in the same direction on comparisons for individual patients were obtained. Genes that concurred as DE among the FTD cells were validated by bootstrapping ANOVA test (k=1000)(43) and with multiplex digital PCR quantification using taqman primers for *RASD2* (Hs04188091), *NEK9* (Hs0092954), *ZFP41* (Hs00905862), *MAPT* (Hs00902194), *GAPDH* (Hs03929097- ThermoFisher) from total RNA samples.

Principal component analyses were performed using normalized CPMs for all post-filtered genes (supplemental Fig. 4A,B) and for genes identified as DE (supplemental Fig. 4C). Cartesian representations for PC1 vs PC2 captured most of the variability as additional principal components represented less than 10% of the expression variability in all cases (supplemental Fig. 4A).

Expression of *MAPT* and *RASD2* were quantified from total RNA (miRvana kit) at day 0 (hiPSCs), day 6 (embryoid bodies), day 12 (neurorossetes), and day 20 (neurospheres) with quantitative PCR using taqman primers specified above. Data was calculated as 2^-ΔCt^ for changes in gene expression level compared to hiPSCs stage. Two-way ANOVA test followed by Tukey HSD post-hoc tests where applicable were used to test differences in means across different cells lines along differentiation stages.

## Acknowledgments

We thank Beverly Davidson for designing and packaging AAV vectors used in this study, and Peter Davies for kindly providing MC1 and PHF-1 antibodies, and Yadong Huang for reprogramming CTR2 and CTR3 iPSCs lines. We also thank the NINDS Cell Repository for the fibroblast lines from the *MAPT* P301L and V337M mutation carriers, as well as Hyung-Seok Kim and Christine Park for reprogramming MAPT-G55R iPSCs Line.

## Funding

This work was funded by the Tau Consortium (K.S.K., A.M.C., C.M.K.), the National Institutes of Health 1U54NS100717 (K.S.K., A.M.C.), R01AG056058 (K.S.K.) and AG046374 (C.M.K.), the National Science Foundation ISS-0808772, ITR 0331697 (S.K.F.), Cohen Veterans Bioscience (K.S.K.) and the Leo and Ann Albert Charitable Trust (K.S.K.). N.J.S. was supported by a postdoctoral EMBO fellowship.

## Author contributions

I.H. and G.L. contributed equally, designing and performing experiments and writing the manuscript. I.H. performed the statistical analyses. J.N.R. performed experiments on Rhes biochemistry. M.G. performed mouse experiments and maintained colonies. D.B. performed immunohistochemistry and microscopy experiments. A.K. performed computational quantifications of brain micrographs and mosaics. C.H. and V.C. performed neuronal differentiation from hiPSCs. E.G. performed IonTorrent sequencing for RNAseq experiments. H.Z. performed RNA sequence mapping and quality control. C.M.K. and A.G. contributed isogenic neurons material for Rhes levels validation and iPS lines generation. C.Z. identified FTD patient with MAPT-G55R, collected skin fibroblasts and prepared primary fibroblast culture. N.J.S., A.D., and A.M.C. designed and performed experiments of lonafarnib effect in proteolysis and lysosomes. S.K.F. performed image analysis and critiqued the manuscript. K.S.K. designed the experiments and wrote the manuscript.

## Note

Supplementary Information and source data files are available in the online version of the paper

---

**Supplemental Figure 1.**
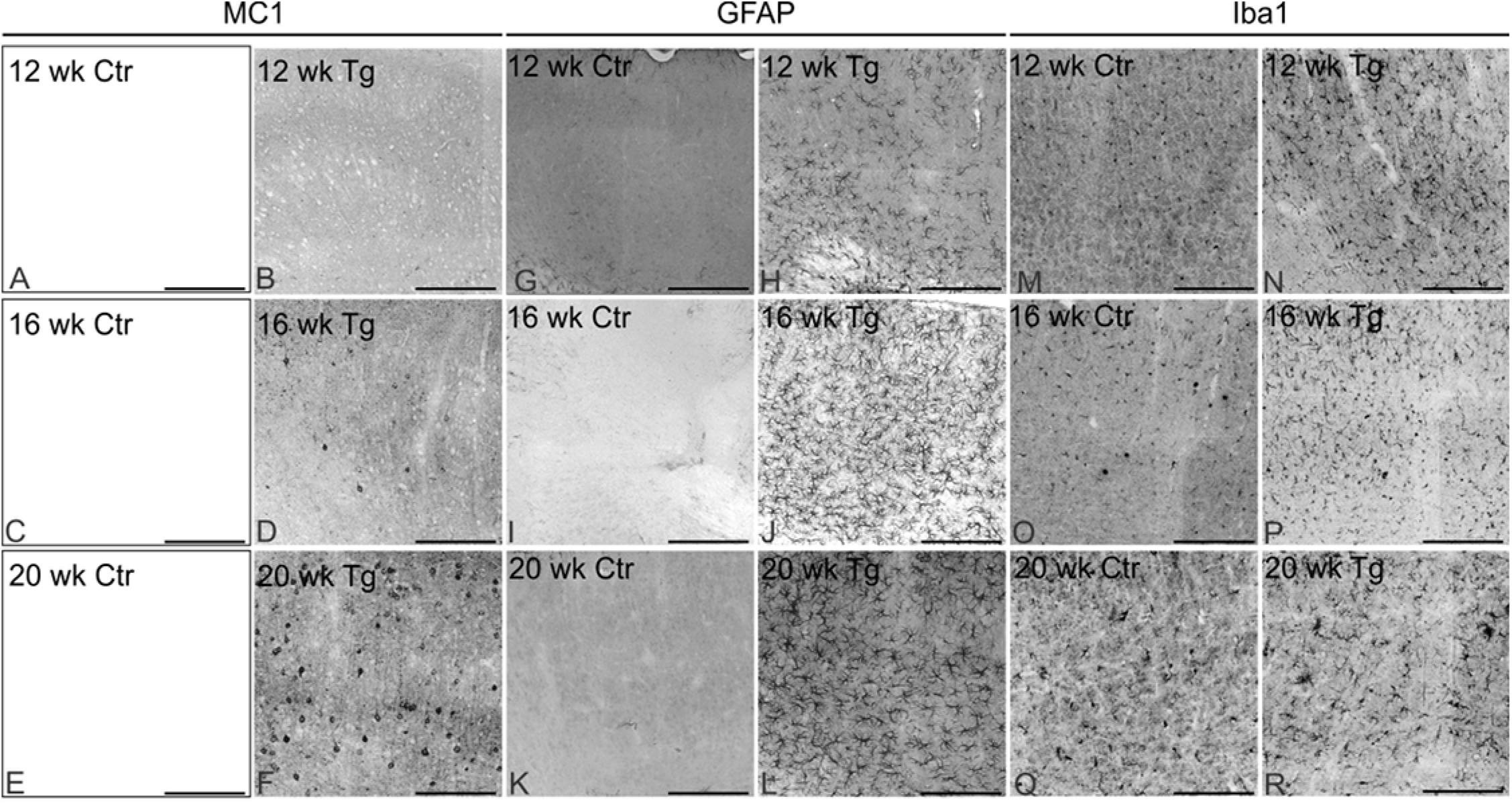
Tau pathology progression in rTg4510 transgenic mice. Cortical micrographs of transgenic rTg4510 (Tg) aged 12 (A-B; G-H; M-N), 16 (C-D; I-J; O-P) or 20 (E-F; K-L; Q-R) weeks-old, displaying the progression of tau pathology, showing no reactivity in control (Ctr) littermates (A, C, E) and age-related increase of tau immunoreactivity with MC1 antibody starting at ~16 weeks (B, D, F). Cortical astrocyte reactivity and density also increased in Tg mice starting at ~12 weeks (H, J, L) in contrast to age-matched Ctr (G, I, K). Microglia reactivity and density in cortex of Tg mice also increased in an age-related manner in transgenic mice (N, P, R), in contrast to Ctr (M, O, Q). Scale bar 100 μm.

**Supplemental Figure 2.**
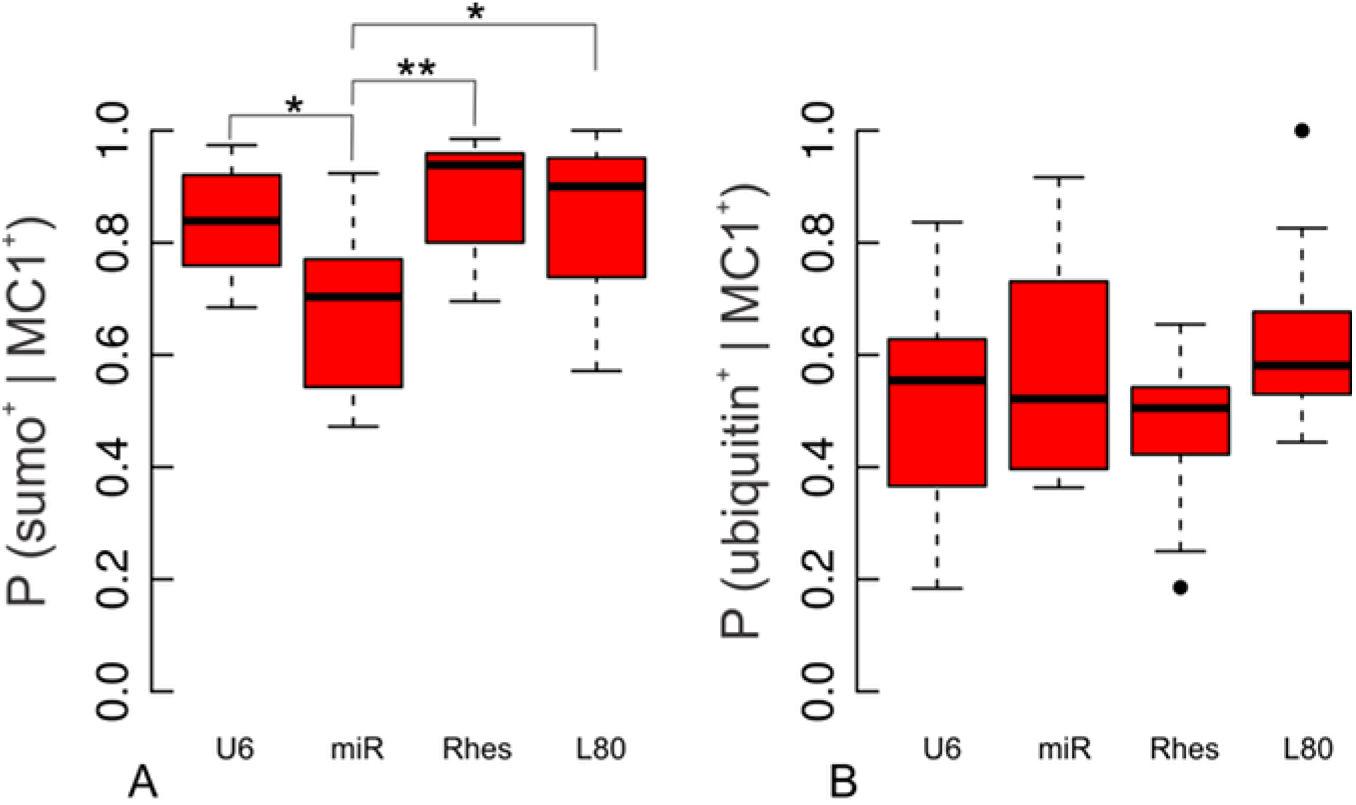
Probability of MC1^+^ cells to label with sumo+ or ubiquitin is not altered by lonafarnib. Cortical micrographs showing double immunostaining using anti-sumo and MC1, or anti-ubiquitin and MC1 (Fig. 5) were used to compute Bayesian probabilities of any cell to be (A) sumo^+^ or (B) ubiquitin^+^ given tau immunoreactivity (MC1^+^). Lonafarnib had no statistically significant effect on either of these probabilities, whereas Rhes-miR altered sumo double-labelling of MC1^+^ cells (sumo^+^ given MC1^+^ ANOVA p=1.61×10^−3^, ubiquitin^+^ given MC1^+^ ANOVA p=0.174). Statistics shown for post-hoc Tukey-HSD test at *p<0.05, **p<0.01.

**Supplemental Figure 3.**
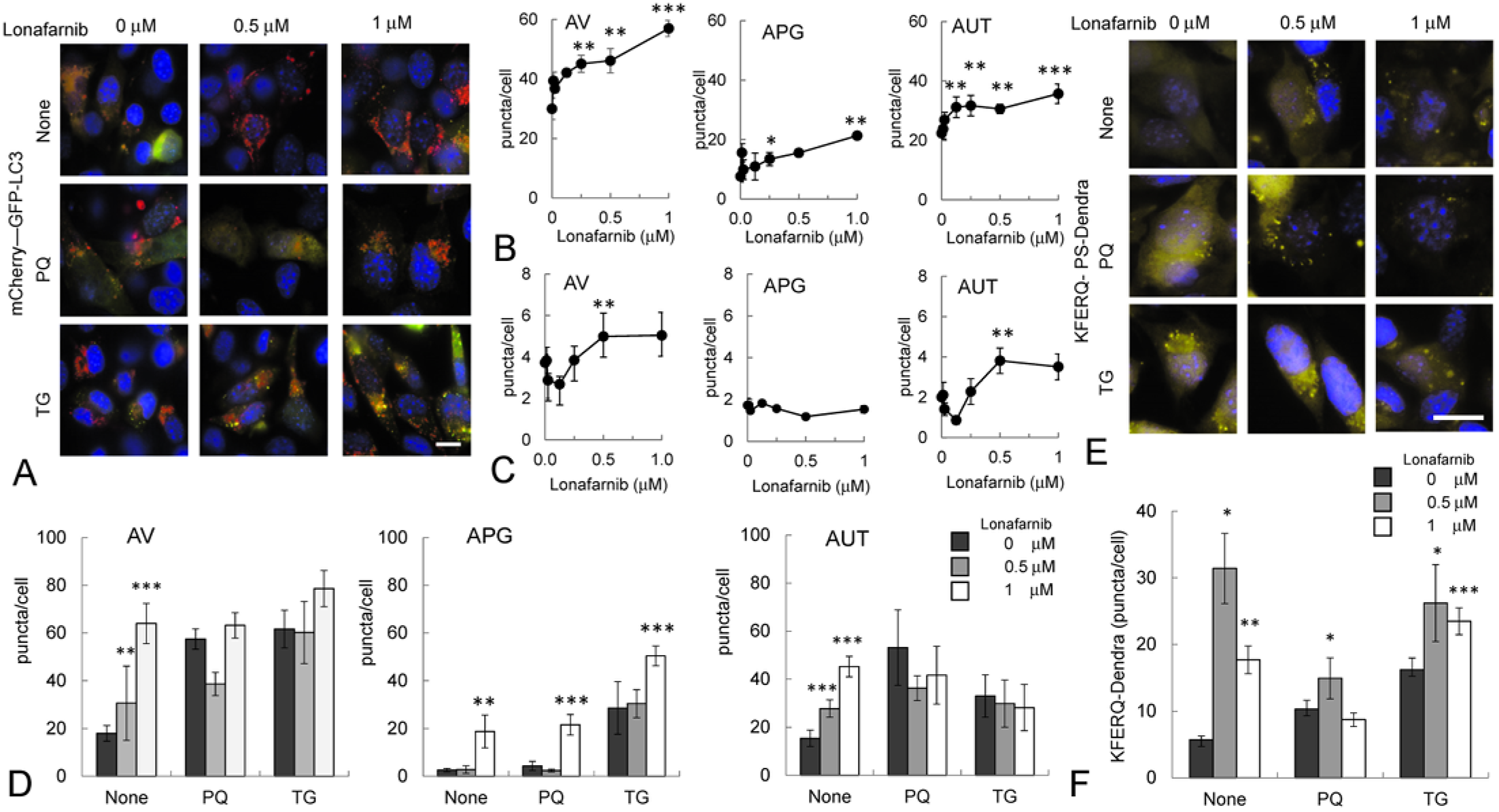
Effect of lonafarnib in macroautophagy and chaperone-mediated autophagy. (A) Representative images of mouse fibroblast in culture (NIH3T3 cells) expressing the tandem reporter mCherry-GFP-LC3B used to quantify the effect of lonafarnib in basal and inducible macroautophagy. Cells were exposed to the indicated concentrations of lonafarnib for 24 h and an additional 24 h in the presence of either paraquat (PQ, 20 μM), or thapsigargin (TG, 300 nM), or left untreated (None). Nuclei are highlighted with DAPI. Dose-dependent effect of treatment with lonafarnib for 48 h of (B) NIH3T3 cells or (C) N2a neuroblastoma cells. Quantified number of autophagic vacuoles (AV), autophagosomes (APG) and autolysosomes (AUT) is shown. (D) Quantification of number of AV, APG and AUT in NIH3T3 cells exposed to the stressors indicated in A (PQ and TG) or left untreated (None). (E) Representative images of NIH3T3 cells expressing the KFERQ-Dendra reporter used to quantify the effect of lonafarnib in basal and inducible chaperone-mediated autophagy (CMA). Cells were photoswitched, exposed to the indicated concentrations of lonafarnib for 24 h and an additional 24 h in the presence of either paraquat (PQ, 20 μM), or thapsigargin (TG, 300 nM), or left untreated (None). Nuclei are highlighted with DAPI. (F) Quantification of the number of puncta positive for photoconverted dendra per cell in cells treated as in E. Quantifications in I-M were done in at least 2,500 cells per condition in three different experiments using high content microscopy. Data is shown as mean ± s.e.m. Differences with untreated (none) are significant for t-test at *p <0.05, **p <0.01 and ***p<1×10^−3^. Scale bar 10 μm.

**Supplemental Figure 4.**
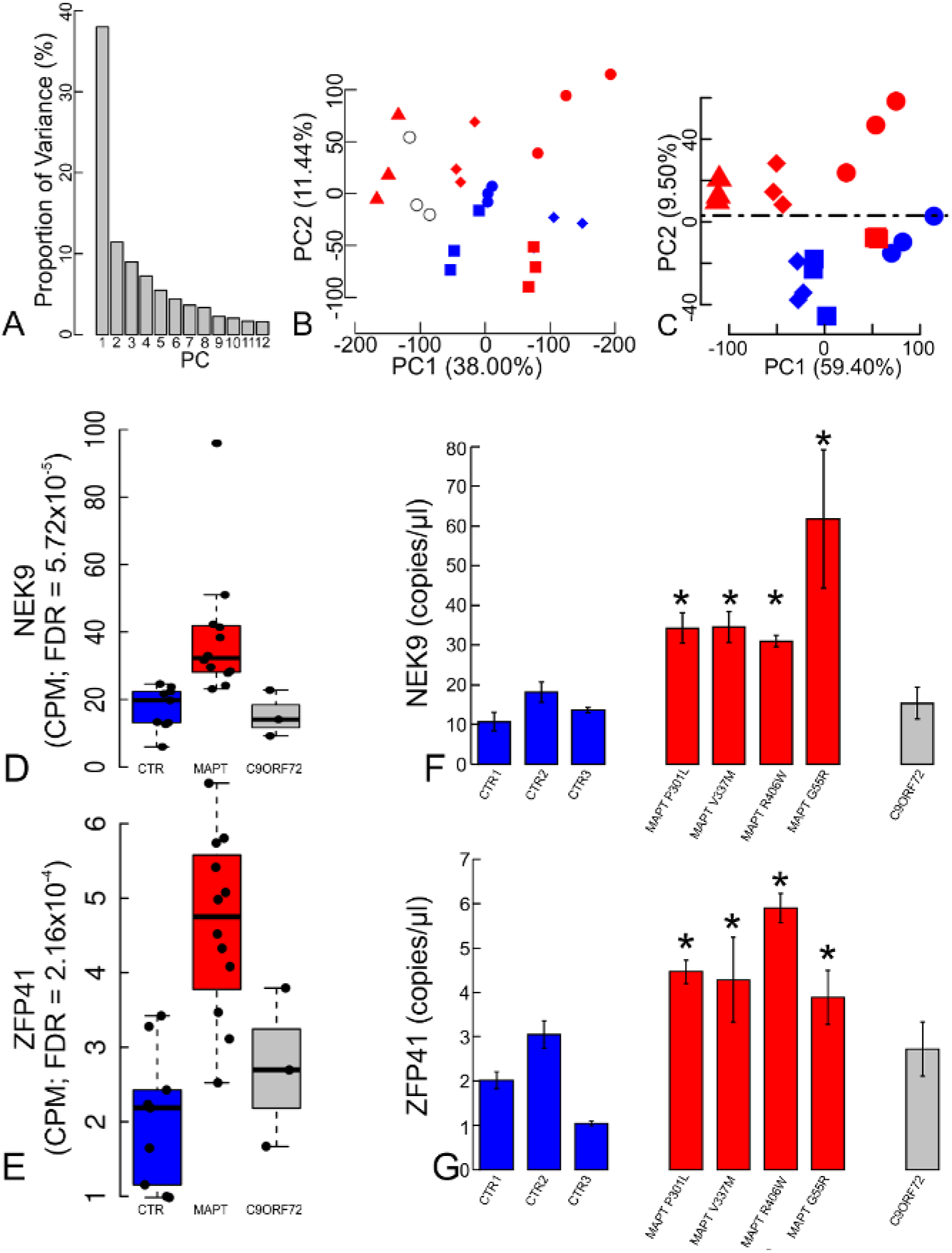
Transcriptomic analysis of hiPSC-derived neurons harboring tau mutations. (A) Principal component analysis (PCA) showing the relative variance contribution of the first 12 principal components (PCs) as calculated from the normalized RNAseq CPMs (C) PC1 and PC2 indicate that individual variation account for most of the RNAseq variation among all transcripts. (B) PCA using normalized CPM of differentially expressed transcripts after comparing all *MAPT* carriers to healthy controls indicates that PC2 can capture variation associated with *MAPT* mutation carriers. Differential expression (DE) analysis compared each *MAPT* mutant hiPSC-derived neuronal line independently to the collective set of controls, revealing between 1746 and 10210 DE genes with a FDR<0.05. Each of these independent comparisons shared at least 7.34% DE genes in common, when intersected pairwise. A Principal Component Analysis (PCA) using CPMs for all expressed genes clustered individual hiPSC lines while triplicates. (D) NEK9 and (E) ZFP41, in addition to RASD2 (Fig 7B) were significantly DE among every tau mutant line assayed. This observation by RNAseq was validated with digital qPCR for (F) NEK9 and (G) ZFP41.

**Supplemental Figure 5.**
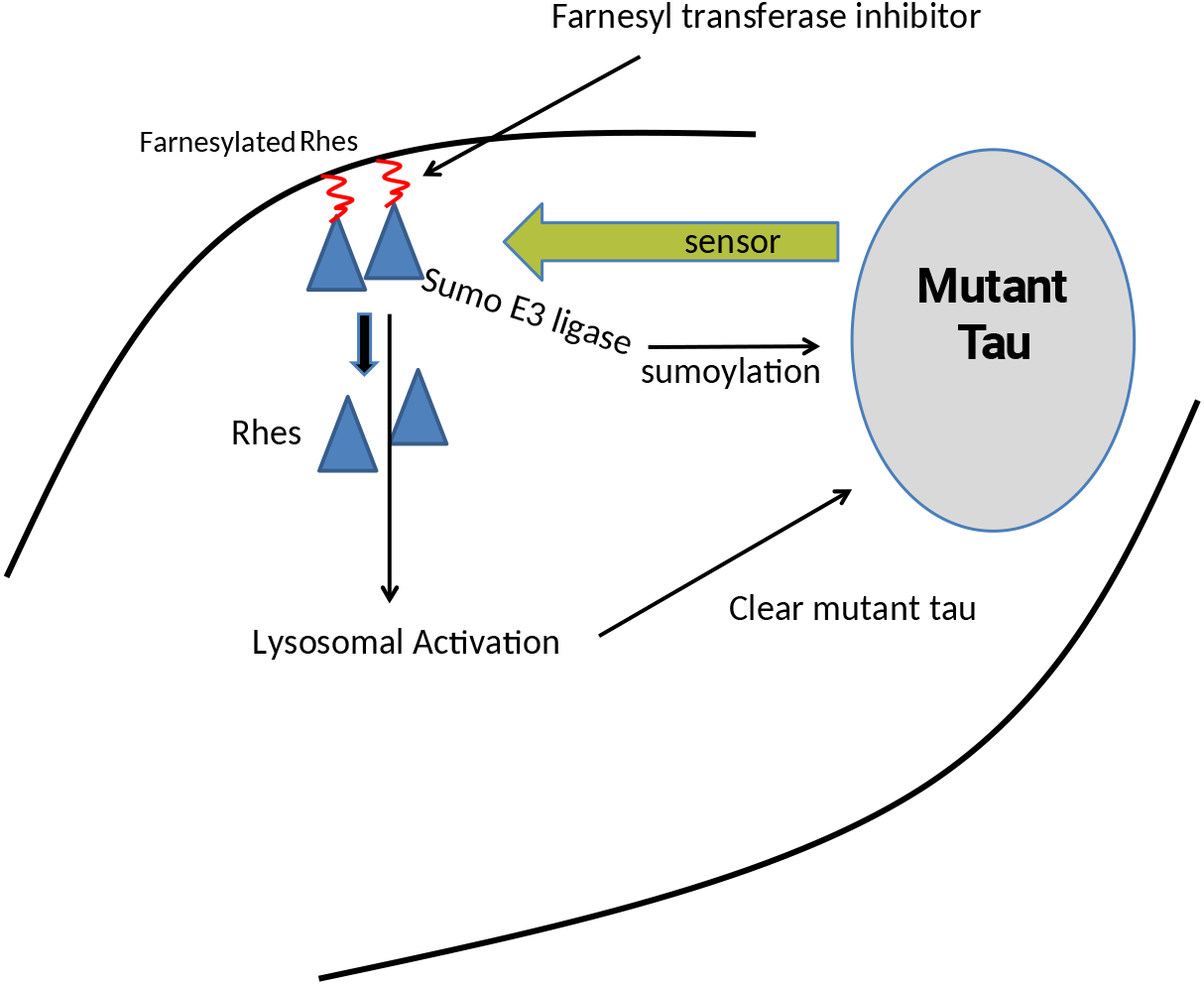
Model for a Rhes pathway mechanism. When mutant tau is present in a cell, a sensor system detects the aberrant protein and releases Rhes from its membrane attachment through a farnesyl group. Inhibiting Rhes farnesylation, which can also be accomplished pharmacologically with a farnesyltransferase inhibitor, results in the activation of the lysosome and clearance of aberrant tau.

## Notes

Competing financial interests: K.S.K. consults for ADRx (Thousand Oaks, CA, U.S.A.)

